# PEDIA-BRAIN: A single nuclei multiomic encyclopedia of the human pons provides a resource for normal development and disease vulnerability

**DOI:** 10.1101/2025.09.21.677597

**Authors:** Tianli Ding, Gary Schweickart, Kaitlin Kaiser, Adithe Rivaldi, Nicholas Marchal, Cole A. Harrington, Aaron Varghese, Ke Qin, Benjamin J. Kelly, Benjamin D. Sunkel, Kathryn L. Stahl, Jack D. Webb, Alex H. Wagner, Jeffrey R. Leonard, Albert M. Isaacs, Katherine E. Miller, Elaine R. Mardis, Michelle A. Wedemeyer

## Abstract

The human pons relays information between the brain and the body. It is affected by pathological processes, including diffuse midline gliomas (DMGs) and multiple sclerosis (MS) which predominantly arise in childhood and middle age, respectively. Although multiple studies address these disease states, a comprehensive resource for normal pons development is lacking. Here we present the first installment of PEDIA-BRAIN, an encyclopedia of gene expression and chromatin accessibility from 140,771 human pons nuclei spanning the first trimester to early adulthood, as a resource for the scientific community. Exploration of the encyclopedia identified two trajectories to mature oligodendrocytes and developmental restriction of genes for neuron to oligodendrocyte progenitor cell synapses. To illustrate the utility of the resource, we compared single cell transcriptomes from DMG and MS tissues to the encyclopedia and identified perturbation of oligodendrocyte subtypes in both diseases. Data may be accessed at https://pediabrain.nchgenomics.org.

## Introduction

The ventral pons is a brainstem structure containing myelinated axon bundles for motor, cerebellar, serotonergic, and noradrenergic arousal systems^1^. Several diseases affect the pons at distinct stages of life, including pontine diffuse midline glioma (DMG), a highly lethal pediatric brain tumor, and multiple sclerosis (MS), which may present with pontine demyelination in adulthood^2, 3^. While morphometric studies of human pons development exist, a cell-type resolved encyclopedia that encompasses disease-relevant periods of development is lacking^4, 5^.

Single nuclei sequencing is accelerating our understanding of the cellular diversity of the human brain. This includes cataloguing of fetal progenitors, regional diversity, inter-individual cell type variation, and studies of disease states. While studies have examined both gene expression and chromatin accessibility, these largely focus on the telencephalon or narrow developmental windows^6–12^. To date, ‘omics studies of the pons have used model organisms or fetal stages for tissue sources^13, 14^.

In general, brain development consists of neurogenesis in utero followed by gliogenesis^8, 15^. While single nuclei sequencing detects oligodendrocyte progenitor/precursors (OPCs) in the hindbrain as early as 5.5 weeks post conception, myelination is a postnatal process that extends into the 3^rd^ decade and drives a six-fold increase in pons volume by 5 years^4, 16–20^. DMGs arise from a missense mutation in histone H3 that disrupts epigenetic regulation of gene expression^21^. Although the susceptibility of the pons to DMGs in childhood indicates that postnatal pons development involves chromatin remodeling, little is known about pons development after age 5.

The ventral pons is densely populated by oligodendrocytes that form insulating myelin wraps around axons. Myelin enables saltatory conduction of axon potentials and provide metabolic support^22^. While postnatal neurogenesis is limited, oligodendrocytes are produced throughout life and have an extraordinarily long life span^23^. Learning of skills throughout life requires remodeling of chromatin and neuron-directed plasticity of synapses and myelin^20, 24^. While the incidence of pontine DMGs decreases precipitously after age 7, the peak onset of multiple sclerosis, an autoimmune disease that leads to demyelination, is in early adulthood^2, 25^. Although both diseases infrequently present outside these windows, their predilection for specific age groups implicates developmentally timed processes in disease susceptibility.

To establish a public resource encompassing the complete spectrum of human pons development, single nuclei RNA sequencing and open chromatin profiling (snRNA-seq + snATAC-seq, or “multiome”) were performed using normal pons tissues corresponding to four age groups: infants and toddlers (0-2 years), children (4-8 years), adolescents (13-14 years), and early adults (31-33 years, i.e. near completion of myelination). Our dataset was integrated with public multiome data from the fetal hindbrain to generate an encyclopedia of gene expression, chromatin dynamics, and cell-cell communications using 140,771 human pons nuclei from the first trimester to the completion of myelination^16^. Data from the resulting PEDIA-BRAIN (**P**ediatric **E**ncyclopedia of **D**evelopment from **I**nfancy to **A**dulthood - **B**rain **R**egions and **A**ssociated **I**ntegrative **N**euroscience) have been deposited at pediabrain.nchgenomics.org as a community resource.

Our analyses identified two convergent trajectories to generate oligodendrocytes, aging signatures of astrocyte and oligodendrocyte subtypes, and a critical period of transsynaptic neuron to OPC communication. We further demonstrated the utility of this encyclopedia via comparison of single nuclei data obtained from pontine DMG and MS disease-affected tissues, revealing a block to oligodendrocyte maturation in the former and oligodendrocyte subtype depletion in the latter^26, 27^. These data solidify the broader utility of developmental encyclopedias for insights into human disease.

## Results

### Core postnatal transcriptional and chromatin accessibility programs of the ventral pons are established at birth

While the dorsal pontine tegmentum contains mixed neuronal populations, the ventral basis pontis contains white matter tracts with few interposed neuron cell bodies. To capture the developmental trajectory of canonical cell types in human pons white matter, paired single nuclei RNA-sequencing and chromatin accessibility sequencing (RNA-seq + ATAC-seq, scMultiome) were performed on ventral white matter from human donor pons aged 0-33 years (n=4-5/group). To construct an encyclopedia of pons development from conception to adulthood, the postnatal single nucleus data were integrated with publicly accessible single nucleus multiome datasets obtained from human fetal first trimester hindbrain which includes progenitors that generate neurons and glia of the pons and cerebellum (n=5, EGAS00001007472)^16^. After quality control, data from a total of 140,771 single nuclei were included for analysis (Figure 1a, Extended Data Fig. 1a-b). Normalization, merging, dimensionality reduction, and clustering were performed in Seurat and Signac^28–30^. A weighted nearest neighbor (WNN) approach was used to integrate paired transcriptional and chromatin accessibility profiles for each nucleus. Subsequent uniform manifold approximation and projection (UMAP) and clustering identified 40 clusters (Extended Data Fig. 1c).

**Fig. 1.**
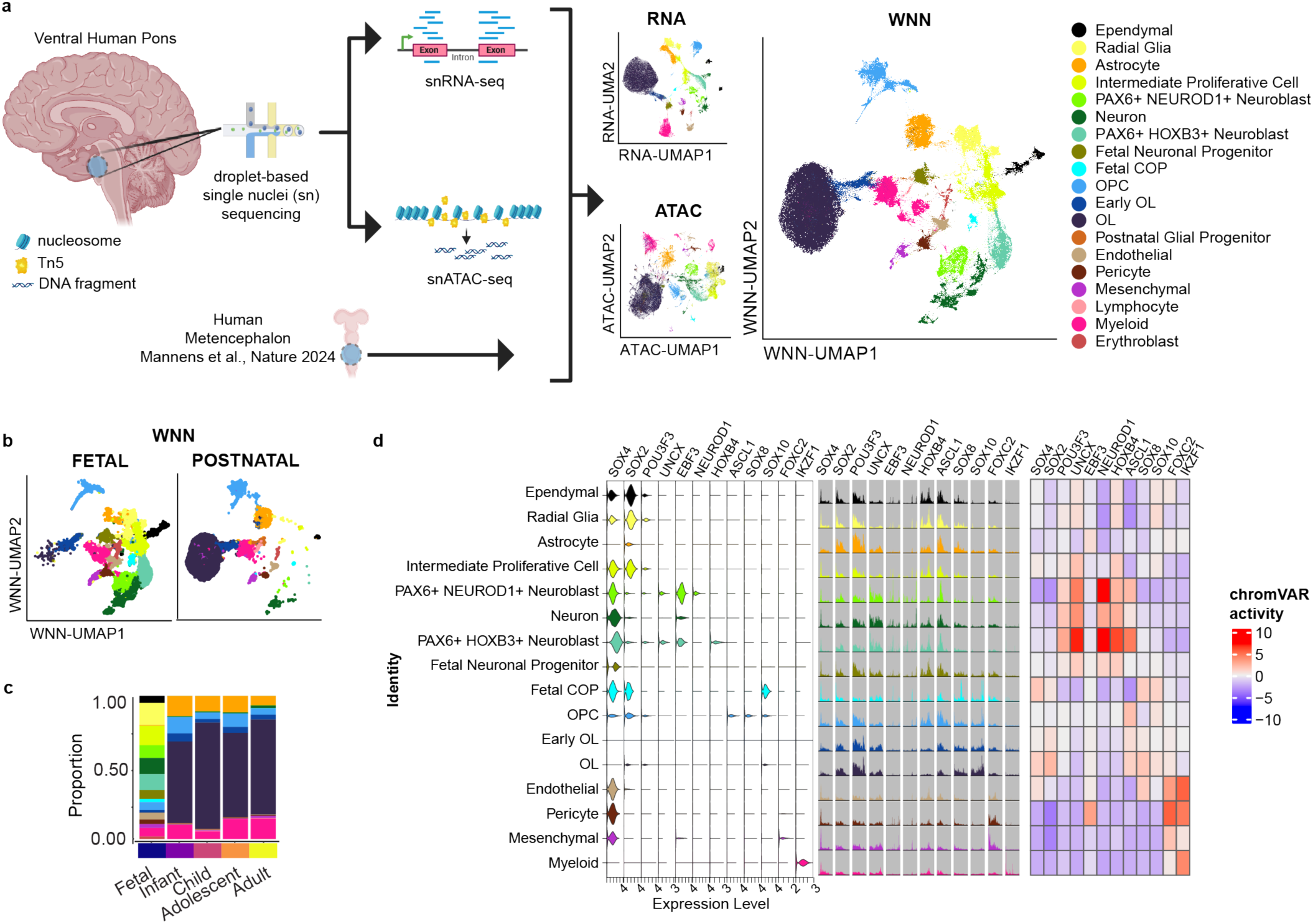
Core postnatal transcriptional and chromatin accessibility programs of the ventral pons are established at birth. Combined single nuclei RNA+ATAC-sequencing identifies transcriptional and chromatin accessibility signatures of cell types in the human pons from the first trimester to early adulthood. **a,** Schematic of experimental design with UMAP of the full encyclopedia of 140,771 nuclei clustered by RNA, ATAC, and WNN with assigned cell types (Created in https://BioRender.com). **b,** WNN UMAP of fetal versus postnatal stages showing transition from fetal progenitors to postnatal glial progenitors and terminally differentiated cell types. **c,** Bar plot of assigned cell type proportions for nuclei from first trimester fetal (n=5; 55,355 nuclei), infant and toddler (n=5; 18,653 nuclei), child (n=5; 25,929 nuclei), adolescent (n=4; 21,493 nuclei), and adult stages (n=4; 19,341 nuclei). **d,** Gene expression, chromatin accessibility, and motif activity of selected cell type specific and regional identify transcription factors for clusters with >1000 cells. Motif enrichment was performed on the top 5000 differentially accessible peaks in each cluster using motifs for TFs expressed in >10% of nuclei in at least one cluster. Motifs with abs(log2(fold change))>1.5 relative to background and BH-corrected p-value<0.05 were considered enriched. Left: Violin plot of gene expression. Middle: chromatin accessibility at the gene body of transcription factors. Right: chromVar motif activity scores for each transcription factor. Abbreviations: UMAP: uniform manifold approximation and projection, ATAC: assay for transposase-accessible chromatin with sequencing, WNN: weighted nearest neighbor, OPC: oligodendrocyte progenitor/precursor cell, COP: committed oligodendrocyte precursor/progenitor cell, OL: oligodendrocyte.

Cluster classification based on canonical markers identified 19 cell types: Ependymal (*PIFO*, *FOXJ1*, *DNAH7*), Radial Glia (*SOX2*, *YAP1*, *FABP7*, *TNC*), Astrocyte (*AQP4*, *ALDH1L1*), Intermediate Proliferative Cell (*CENPF*, *TOP2A*, *PAX3*), PAX6+ NEUROD1+ Neuroblast (*DCX*, *PAX6, PAX3*), Neuron (*MAP2*, *TUBB3*, *RBFOX3*), PAX6+ HOXB3+ Neuroblast (*DCX*, *PAX6*, *HOXB3*), Fetal Neuronal Progenitor (*STMN2*, *TUBB3*), Fetal Committed Oligodendrocyte Progenitor (COP: *SOX2*, *YAP1*, *SOX10*, *TCF7L2*), Oligodendrocyte Progenitor (OPC: *PDGFRA*, *OLIG2*), Early Oligodendrocyte (OL: *PLP1*, *MBP*), Oligodendrocyte (OL: *MBP*, *CNP*, *MOBP*), Postnatal Glial Progenitor (*GFAP*, *MOBP*), Endothelial (*PECAM1*), Pericyte (*PDGFRB*), Mesenchymal (*LUM*), Lymphocyte (*CD3D*, *PTPRC*), Myeloid (*PTPRC*, *P2RY12*), and Erythroblast (*GYPA*) (Extended Data Fig. 1d). Gene expression in cluster 11 was driven by high expression of lymphocyte genes in a small number of cells. Subsequent subclustering resolved one cluster of lymphocytes, one cluster of postnatal glia progenitors, and 6 clusters of fetal neuronal progenitors (Extended Data Fig. 1e-f). Cell type assignments were strongly weighted by both RNA and ATAC signatures (Extended Data Fig. 2a). By infancy, the gene expression and chromatin accessibility signatures of canonical postnatal CNS cell types were established and readily distinguished from fetal progenitors. Few glial cells were identified in fetal samples consistent with neurogenesis preceding gliogenesis (Fig. 1b-c, Extended Data 2b, Supplementary Table 1-3).

Differential gene expression, motif enrichment of differentially accessible peaks, and chromVar transcription factor (TF) activity scores were calculated for all cell types with >1000 cells (Fig. 1d, Extended Data Fig. 2d, Supplementary Table 1-3). Although chromatin accessibility was retained postnatally around TFs with roles in neural development (*ASCL1*, *SOX4*, *SOX2*, *POU3F3*) and developmental patterning of the pons (*HOXB4*), both TF expression and chromVar activity scores revealed more cell type specific roles (Fig. 1d). Both expression and chromVar activity of neuronal TFs *UNCX*, *NEUROD1*, and *HOXB4* were highly specific to neurons. *SOX10* was specific to oligodendrocyte lineage cells while *FOXC2* and *IKZF1* were specific to non-neuronal cell types.

Although the ventral pons is predominantly white matter, a small population of neurons was identified in all postnatal age groups. Given the predominance of neuronal progenitors in fetal samples, fetal radial glia and progenitors were subclustered with postnatal neurons which identified diverse excitatory (*SLC17A6*) and inhibitory (*GAD1/2*) neuronal progenitor populations and regional transcription factors involved in hindbrain patterning (*PAX6*, *HOXB3*, Extended Data Fig. 3a-e). Several clusters expressed the glycine transporter *SLC6A5* (norepinephrine transporter 1) consistent with glycinergic neurons. Postnatal neurons from the ventral pons represented a mixed population of *RBFOX3+/DCX-*postmitotic neurons that expressed the calcium sensor *VSNL1*, a biomarker of neurodegeneration, and the opiate receptor *OPCML* (Extended Data Fig. 3f)^31^.

### Integrated analysis of gene expression and chromatin accessibility distinguishes astrocyte subtypes

Though radial glia and astrocytes are temporally and morphologically distinct cell types, radial glia have been proposed to transition to astrocytes postnatally^32^. Here, radial glia and astrocyte clusters expressed overlapping markers (*SOX2*, *YAP1*, *TNC*, *MAP2)* and were closely approximated by principal component analysis of gene expression signatures (Fig. 1a, Extended Data Fig. 1d)^33^. Integrated RNA and ATAC signatures (WNN) readily distinguished radial glia from astrocytes and identified 4 subtypes of postnatal astrocytes: *EAAT2-hi*, *CD44-hi*, undetermined astrocytes, and a small cluster of Oligo-like astrocytes that expressed the oligodendrocyte maker *MOBP* (Fig. 2a, Extended Data Fig. 4a-b). Radial glia were enriched for biologic processes involved in development, *CD44-hi* astrocytes were enriched for processes involved in neuron projection development, and *EAAT2-hi* astrocytes were enriched for processes involved in behavior and transsynaptic signaling (Fig. 2b, Supplementary Table 4). While no significant biologic processes distinguished Undetermined astrocytes from other subtypes, principal component analysis of gene expression was suggestive of a transitional state between radial glia and postnatal astrocytes (Extended Data Fig. 4b). *EAAT2-hi* astrocytes were enriched for the glutamate re-uptake transporter *SLC1A2/EAAT2* and the WNT regulator *LGR6* while *CD44-hi* astrocytes were enriched for *CD44*, a marker of fibrous white matter astrocytes (Extended Data Fig 4c-d)^34^. Postnatal astrocytes expressed higher levels of *CD44* and *IDH2-DT*, a lncRNA implicated in stroke (Fig. 2c)^35^. *CD44-hi* astrocytes were strongly enriched for the *ROBO2* receptor which mediates axon guidance in the developing hindbrain (Fig. 2d)^36^.

**Fig. 2.**
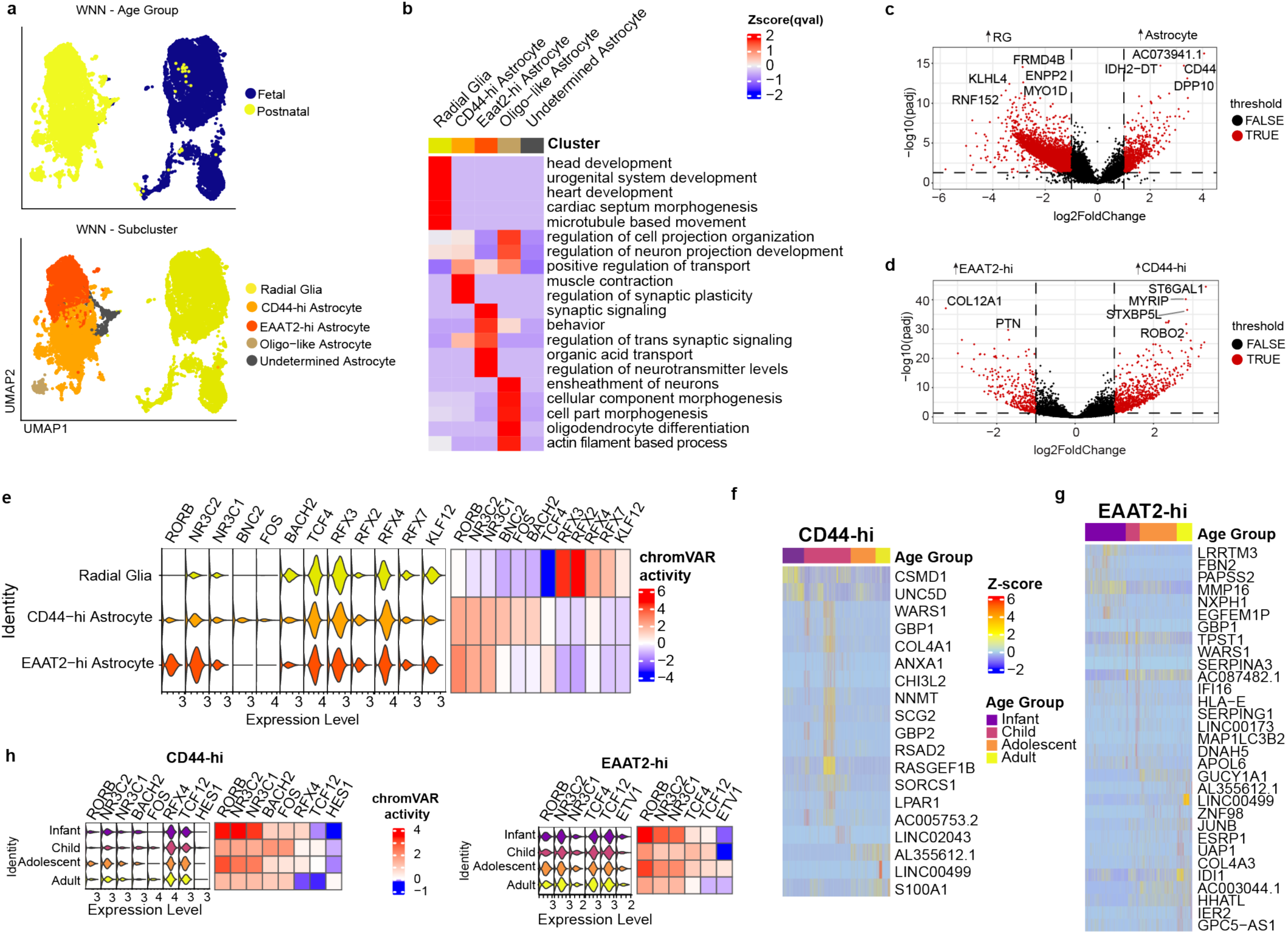
Integrated gene expression and chromatin accessibility distinguishes astrocyte subtypes. **a,** WNN UMAP of subclustered radial glia and astrocytes grouped by developmental stage (Top) and assigned cell type (Bottom) showing clear separation between fetal radial glia and postnatal astrocytes. Postnatal astrocytes separated into *EAAT2-hi* and *CD44-hi* subtypes with smaller clusters of Oligo-like and Undetermined astrocytes. **b,** Heatmap of normalized Q-values for the top 5 Gene Ontology Biologic Processes for each subtype. Gene ontology was performed using differentially upregulated genes from each cluster defined by abs(log_2_(fold change))>1.5 and BH-corrected p-value<0.05 relative to other cell types. **c-d,** Volcano plot of differentially expressed transcripts between postnatal astrocytes and radial glia (**c**) and between *CD44-hi* and *EAAT2-hi* astrocytes (**d**). Red dots denote genes with BH-corrected p-value<0.05 and abs(log_2_(fold change))>1.5. **e,** Violin plot of transcript expression (Left) and heatmap of chromVar motif activity (Right) for transcription factors with significant motif enrichment in radial glia and astrocyte subtypes. chromVar scores represent values z-scored by individual nucleus across the full encyclopedia. Enriched motifs were identified as described for Fig. 1 then filtered for mean RNA counts>1.5 and mean chromVar activity>0.1. **f-g,** Heatmap of normalized transcript expression for genes that are differentially expressed during postnatal development of *CD44-hi* (**f**) and *EAAT2-hi* (**g**) astrocytes as defined by expression in >5% of cells, log(fold change)>1, and BH-corrected p-value<0.05 in at least one age group. **h,** Violin plot of transcript expression (Left) and corresponding chromVar motif activity (Right) for transcription factors with significant motif enrichment in *CD44-hi* and *EAAT2-hi* astrocytes.

Given the significant overlap in gene expression between radial glia and astrocytes, we examined TF gene expression, motif enrichment, and chromVar activity to uncover putative TF regulators that drive their divergent behavior (Fig. 2e). While radial glia were enriched for regulatory factor X transcription factors (*RFX2/3/4*), astrocytes were enriched for nuclear hormone receptors (*RORB*, *NR3C2*, *NR3C1*) which encode retinoic acid, glucocorticoid, and mineralocorticoid receptors, underscoring the homeostatic function of astrocytes. Both radial glia and astrocytes expressed RFX factor transcripts and depletion of Tn5 insertion sites around RFX factor motifs consistent with protection from bound TFs. In contrast, chromVar motif activity and overall accessibility were significantly enriched around RFX factor genes in radial glia relative to astrocytes suggestive of increased chromatin accessibility for RFX-regulated genomic loci in radial glia (Fig. 2e, Extended Data Fig. 4e, Supplementary Table 5). *CD44-hi* astrocytes were further enriched for peaks regulated by the immediate early gene *FOS*, the zinc finger protein *BCN2*, and the basic leucine zipper TF *BACH2*.

Astrocytes regulate synapse formation, pruning, and maintenance. As age-related declines in neuroplasticity have been partially attributed to astrocytes, gene expression and TF motif enrichment were investigated across postnatal development. *CD44-hi* astrocytes expressed the axon guidance gene *UNC5D*, during infancy, the synaptic regulator *SORCS1* during childhood, and S100A1, a chaperone protein associated with the Hsp70/Hsp90 chaperone complex during adulthood (Fig 2f, Supplementary Table 6)^37, 38^. *EAAT2-hi* astrocytes were notable for expression of *LRRTM3*, a regulator of excitatory synapse development, during infancy and of *HHATL*, an inhibitor of SHH signaling during adulthood (Fig. 2g)^39^. Both major subtypes of astrocytes exhibited periods of increased chromVar activity for *RORB*, *NR3C2*, *NR3C1* during infancy and adolescence. Notably, activity for *BACH2* and the immediate early gene *FOS* were specifically enriched in CD44-hi astrocytes during childhood (Fig 2h, Supplementary Table 6). TF footprinting for *FOS* and *BACH2* identified a dip in the expected Tn5 enrichment around *FOS* and *BACH2* motif binding sites at all ages, suggesting protection from bound TFs, however, elevated Tn5 insertion frequency in the flank regions during childhood was suggestive of increased accessibility during this period (Extended Data Fig. 4f). These data indicate that changes in astrocyte function during development may be regulated by specific transcription factor modules.

### Convergent developmental programs give rise to mature oligodendrocytes

As the ventral pons contains white matter tracts that undergo myelination during infancy and childhood, the developmental trajectories and aging signatures of oligodendrocyte lineage cells were explored. Two distinct clusters of progenitor cells expressed the transcription factor *SOX10* which activates myelin programs in both the peripheral and central nervous system, suggestive of two developmental trajectories for the generation of oligodendrocytes (Fig 1a,d)^40^. Fetal committed progenitors (fetal COP) are distinguished by high expression of *SOX2* and the TEAD binding partner *YAP1* while a separate cluster of fetal and postnatal cells cluster with postnatal OPCs and express classical OPC markers (*OLIG2*, *PDGFRA*, Extended Data Fig. 5a-c). To explore the trajectories of cell types committed to the oligodendrocyte lineage, *SOX10+* progenitors were subclustered together (early oligodendrocytes (Early OL) and myelinating oligodendrocytes (OL, Fig. 3a-c)). Both progenitor populations converged on a *PLP1+* early oligodendrocyte stage (Fig. 3d). A declining number (nCount) and diversity (nFeature) of both RNA transcripts and accessible ATAC peaks was noted from COP to OPC with a nadir in early oligodendrocytes and a burst of transcription in mature OLs (Fig. 3e).

**Fig. 3.**
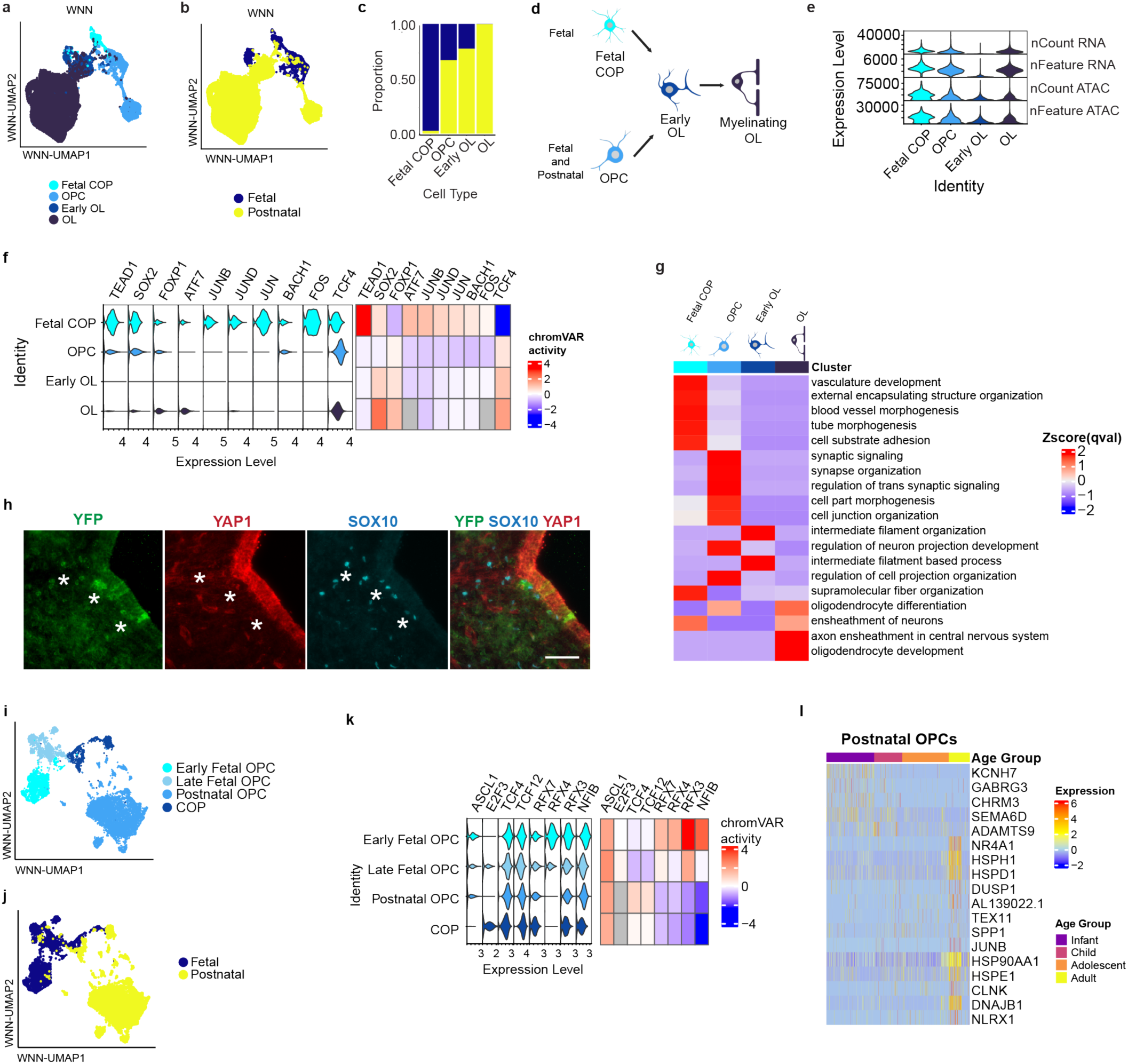
Convergent developmental programs give rise to mature oligodendrocytes. **a-b,** WNN UMAP of subclustered oligodendrocyte lineage cells including fetal COP, OPC, early OL, and OL from full encyclopedia grouped by assigned cell type (**a**) and tissue of origin (**b**). **c,** Barplot showing proportional of nuclei from fetal vs postnatal tissues in each subgroup. **d,** Schematic of proposed developmental trajectories for fetal-restricted and classical fetal/postnatal oligodendrocyte trajectories (Created in https://BioRender.com). **e,** Stacked violin plot of nCount and nFeature for RNA and ATAC reads by cell type shows reduced RNA counts and features at the early OL stage. **f,** Violin plot of transcript expression (Left) and chromVar motif activity (Right) for transcription factors with significant motif enrichment in oligodendrocyte lineage subtypes. Motif enrichment was defined as in Fig. 2. **g,** Heatmap of normalized Q-values for the top 5 Gene Ontology Biologic Processes for each cell type. Gene ontology was performed using differentially upregulated genes from each cluster defined by abs(log_2_(fold change))>1.5 and BH-corrected p-value<0.05 for each OL lineage cell type relative to other OL lineage cell types. **h,** Immunofluorescent images of the ventromedial 4^th^ ventricle ependymal region of E18.5 *Sox10-Cre;ROSA26-YFP* mice. Co-localization of YFP+ (green), YAP1 (red), and SOX10+ nuclei (cyan) indicates SOX10-Cre mediated activation of YFP in SOX10+/YAP1+ cells of the subventricular zone. Scale bar: 50μm. **i-j,** WNN UMAP of subclustered OPCs from full encyclopedia grouped by assigned cell type (**i**) and tissue of origin (**j**). **k,** Violin plot of transcript expression (Left) and corresponding chromVar motif activity (Right) for transcription factors with significant motif enrichment in OPC subtypes. **l,** Heatmap of normalized expression for genes that are upregulated in OPCs in at least one age group with log(fold change)>1 and BH-corrected p-value<0.05. Abbreviations: nCount: total number of reads, nFeature: total number of reads corresponding to unique features (genes/peaks).

Given the paucity of differentially expressed genes or differentially accessible peaks in early OLs relative to other OL lineage clusters, motif enrichment of expressed transcription factors was investigated. Fetal COPs showed both higher expression and increased chromVar activity scores for the YAP1 binding partner *TEAD1*, the neural stem cell marker *SOX2*, and the AP-1 complex transcription factors (*FOS*, *JUN*, *JUNB*, *JUND*) while OPCs, early OLs, and OLs were enriched for peaks with the *TCF4* motif (Fig. 3f, Extended Data Fig. 5d, Supplementary Table 5). Gene ontology analysis was performed to identify biologic processes that distinguished each cluster (Fig. 3g, Supplementary Data Table 4). Fetal COPs were enriched for external encapsulating structure organization and vasculature development. Concordant with reports that synaptic communication from neurons regulate OPC behavior, OPCs were highly enriched for synaptic signaling and synapse organization^41^. Given restricted access to human fetal tissues, we sought to support these findings in a mouse model. Here, Sox10-Cre mice were crossed with ROSA26-LSL-YFP mice and euthanized at E18.5^42^. Identification of YFP+/SOX10+/YAP1+ progenitors in the ventromedial 4^th^ ventricle subependymal region of the mouse pons was suggestive of the presence of analogous *YAP1+/SOX10+* committed oligodendrocyte progenitors in the embryonic mouse pons (Fig 3h).

While the *YAP1+/SOX10+* fetal COP cluster was restricted to fetal stages, PDGFRA+ OPCs are identified at all developmental stages. Myelination is a postnatal process mediated by differentiation of PDGFRA+ OPCs into mature oligodendrocytes. Although initial pons myelination occurs during infancy, PDGFRA+ OPCs are recruited by active neuronal circuits to generate new myelinating oligodendrocytes throughout life and generation of new myelin is necessary for learning of new tasks^4, 41^. To understand the decline in myelinating potential with aging, OPCs were subclustered. While WNN readily identifies fetal and postnatal OPC clusters, both states converge on a GPR17+/MYRF+ committed oligodendrocyte progenitor state (COP, Fig. 3i-j). Although *ASCL1* chromVar activity scores were elevated in all OPC subtypes, fetal OPCs were enriched for *RFX3/4/7* and *NFIB* motifs while postnatal OPCs were enriched for *TCF4/12* motifs (Fig 3k, Supplementary Table 5). At the transcript level, fetal OPCs were enriched for markers of proliferation (*TOP2A*, *MKI67*, Extended Data Fig. 5e). The observation that multiple TF modules can give rise to cell types with an OPC gene expression signature is concordant with histologic studies in mice identifying a perinatal transition from SOX2/PDGFRA+ OPCs to the classical OLIG2+/PDGFRA+ OPC^43^.

Although motif signatures were stable across postnatal development in OPCs, aging was associated with a shift from genes related to synaptic communication and axon guidance (*KCNH7*, *GABRG3*, *CHRM3*, *SEMA6D*) to genes associated with stress responses (*HSPD1/E1/H1*, *SPP1*) (Fig. 3i, Supplementary Table 6). Thus, adult OPCs maintain a stable core transcriptional program but gradually acquire features of senescence that may impede their capacity to produce mature OLs.

### Oligodendrocytes accumulate markers of senescence by early adulthood

Subclustering of postnatal early oligodendrocytes (early OL) and oligodendrocytes (OL) identified a continuum of transcriptional states that was present in all postnatal age groups (Fig. 4a-c). Pseudotime trajectory analysis indicated a transition from *NKX6-2-*early OL to *NKX6-2+* OL. *NKX6-2*+ mature OLs undergo a shift from *OPALIN+/NLGN1+* OL to *RBFOX+/S100A1+* OL (Fig 4d-f). *OPALIN*+ OLs were enriched for genes involved in mitosis, chromatin organization, and histone modification while *RBFOX1*+ OLs were enriched for genes involved in metabolic processes necessary for saltatory conduction (Fig. 4g-h). As the larger size and limited availability of human donor pons precluded spatial characterization of the human ventral pons, axial sections of the pons from P8 C57BL6 mice were stained for SOX10, Opalin, and RBFOX1 to explore whether *RBFOX1+* vs *Opalin+* OLs represent regional subtypes versus transitional stages of OL development (Fig. 4i). As expected, Opalin, which localizes the myelin paranodes and declines with aging, was ubiquitously expressed in myelin processes and perinuclear staining for Opalin was identified in the majority of SOX10+ OL^44^. While spatially restricted RBFOX1 vs Opalin OLs were not identified, an interspersed subpopulation of SOX10+/RBFOX1+ OLs was identified that lacked perinuclear Opalin and may represent a subpopulation of OLs that is not actively forming new myelin wraps. Subsequent immunostaining of human ventral pons from an adult donor confirmed the perinuclear localization of the Hsp70/Hsp90 chaperone protein S100A1 in SOX10+/MBP+ oligodendrocytes and may represent a marker of stress or senescence in OLs (Fig. 4j). While Opalin OLs expressed glutamate receptors (*GRM8*, *GRIK2*), the cytoskeletal gene *SGCZ*, and the paranode gene *CNTN1* during infancy, adult *Opalin+* OLs expressed higher levels of different paranode gene *CNTNAP2/CASPR2*, possibly indicating shifts in paranode composition with development. While *RBFOX1+* OLs also expressed higher levels of *CNTN1* and *SGCZ* during infancy, adult *RBFOX1+* OLs expressed higher levels of the HHATL, an inhibitor of hedgehog signaling, and senescence markers (*CLNK*, *DUXAP10*, *NUPR1*, Fig. 4k, Supplementary Table 6). As hedgehog signaling promotes OPC maturation, the decline in genes associated with nodes of Ranvier and upregulation of genes associated with inhibition of hedgehog signaling (*HHATL*) and senescence (*NUPR1*, *CLNK*, *DUXAP10*) may contribute to the decreased efficiency of remyelination in adults.

**Fig. 4.**
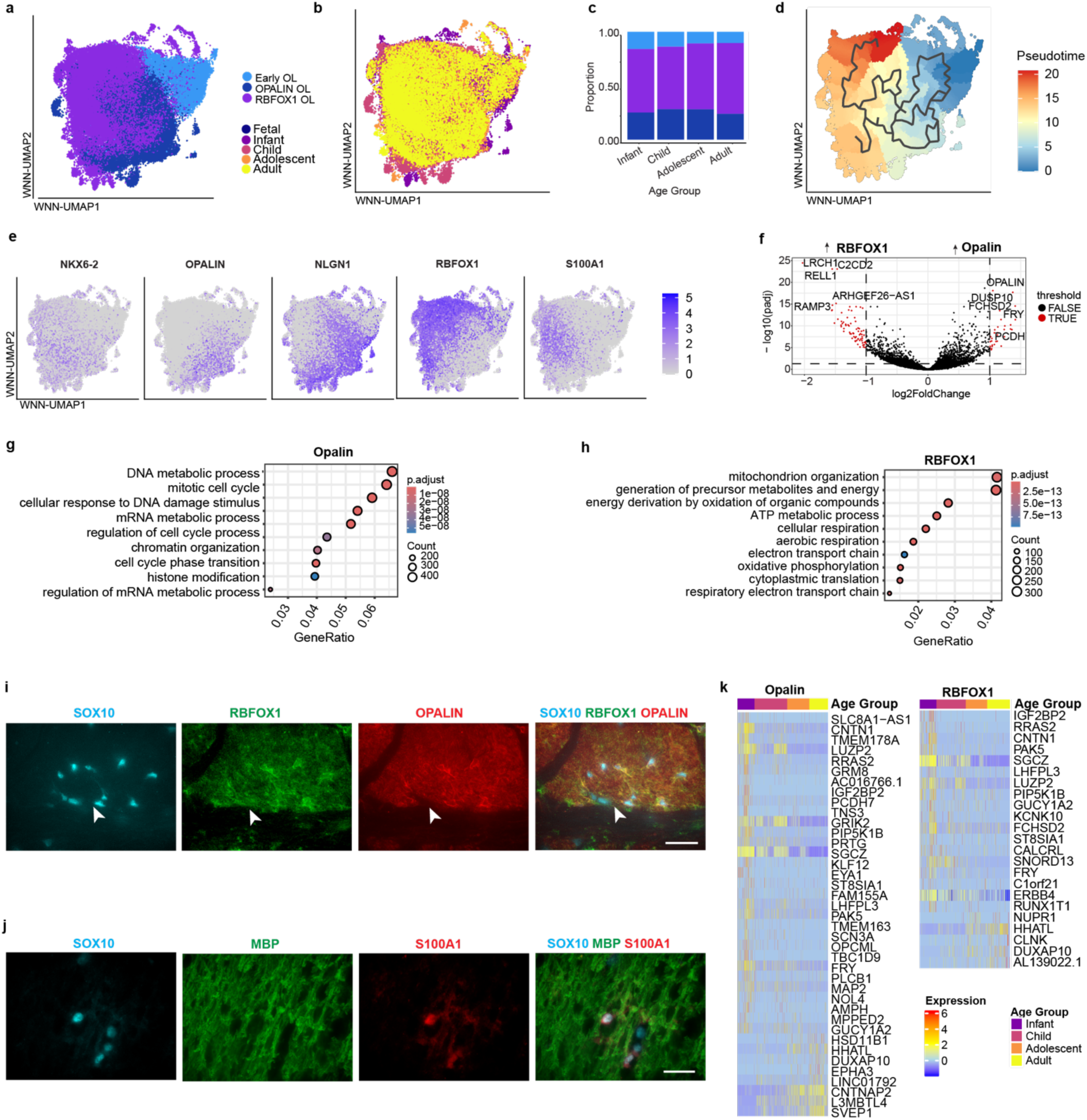
Oligodendrocytes accumulate markers of senescence by early adulthood. **a-b,** WNN UMAP of oligodendrocyte progenitor/precursor lineage cells sub-clustered by gene expression and grouped by subtype (**a**) and age group (**b**). **c,** Barplot showing distribution of oligodendrocyte subtypes across postnatal development. **d,** Pseudotime trajectory analysis of oligodendrocyte lineage cells using early OLs as a starting point. **e,** Feature plots showing markers on the continuum of *Opalin+* to *RBFOX1+* OLs. **f,** Volcano plot of gene expression in *OPALIN+* vs. *RBFOX1+* oligodendrocytes. Red dots denote genes with BH-corrected p-value<0.05 and abs(log_2_(fold change)) >1.5. **g-h,** Dot plot of top 10 gene ontology biological processes enriched in *Opalin+* vs *RBFOX1+* OLs. **i,** Immunostaining of white matter tracts in axial sections of ventral pons of P8 mice identifies SOX10+/RBFOX1+ oligodendrocytes without perinuclear expression of Opalin (white arrow). Scale bar: 20μm. **j,** Immunostaining of ventral white matter from the pons of a 33yo male donor of African descent identifies MBP+/SOX10+ oligodendrocytes with perinuclear expression of S100A1. Scale bar: 20μm. **k,** Heatmap of normalized expression of genes that are upregulated in *Opalin+* (Left) or *RBFOX1+* (right) OLs in at least one age group defined as log(fold change)>1 and BH-corrected p-value<0.05.

### Receptor-ligand communication probabilities identify critical periods for neuroplasticity in the ventral pons

Reasoning that brain development would be accompanied by shifts in cellular communication, CellChat was used to calculate communication probabilities of receptor ligand signaling during postnatal pons development. Top signaling pathways involved trans-synaptic adhesion genes and pathways that have been implicated in either long term potentiation at neuron-neuron synapses or at the neuron-OPC synapses that regulate OPCs (Fig. 5a-b). Cell-cell communication probabilities for *CD6*, myelin protein zero-like (*MPZ*), and *NOTCH* pathways involved in OPC differentiation showed significant shifts across postnatal development with loss of inferred outgoing DLL3-NOTCH1/2/3 communication from OPCs in adulthood (Extended Data Fig. 6, Supplementary Table 7)^45^.

**Fig. 5.**
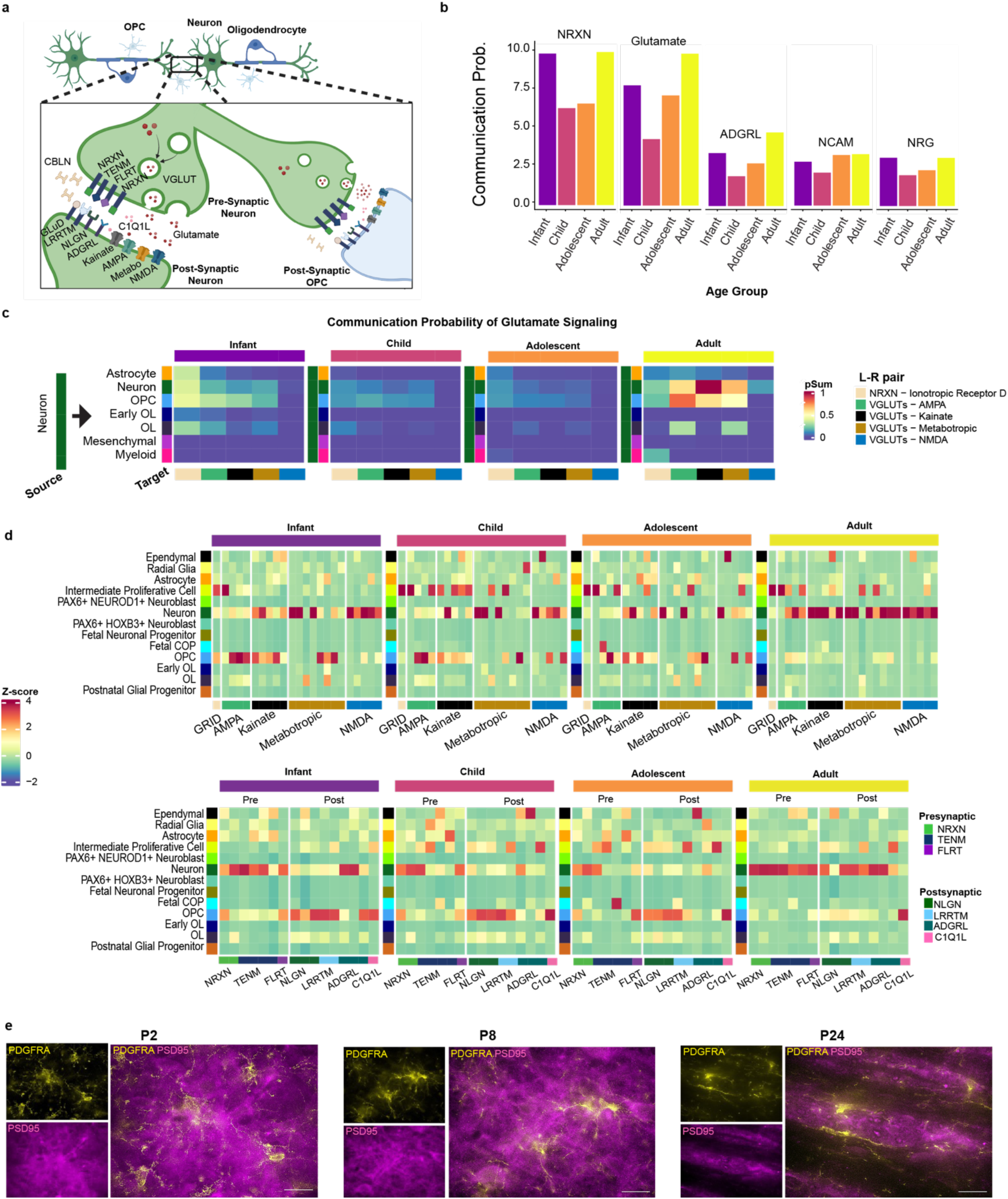
Receptor-ligand communication probabilities identify critical periods for neuroplasticity in the ventral pons. **a,** Schematic of glutamate signaling at neuron-neuron and neuron-OPC synapses (Created in https://BioRender.com). **b,** Bar plot of net communication probabilities for the top 5 receptor-ligand signaling pathways across postnatal pons development. **c,** Heatmap of communication probabilities for glutamate signaling from neurons to surrounding cell types across development. **d,** Heatmap of normalized gene expression of glutamate receptors (Top) and transsynaptic strengthening genes (Bottom) showing robust expression in OPCs during development. **e,** Immunostaining of axial sections of the murine ventral pons white matter identifies PDGFRA+ OPCs (yellow) surrounded by diffuse PSD95+ puncta corresponding to the postsynaptic density of excitatory synapses (magenta) at P2 and P8. By P24, OPCs are aligned with discrete regions of PSD95+ puncta corresponding to the myelinating transverse pontocerebellar white matter tracts. Scale bar: 20μm. Abbreviations: ADGRL: adhesion G protein-coupled receptor L1 (lactrophilin); NRXN: neurexin; NRG: neurogenin; NCAM: neural cell adhesion molecule; Commun. Prob: Communication probability; EAAT: excitatory amino acid transporter; CBLN: cerebellin; VGLUT: vesicular glutamate transporter; GRID: glutamate ionotropic receptor delta type subunit; GRIA: glutamate ionotropic receptor AMPA type; GRIK: glutamate ionotropic receptor kainate type; GRIM: metabotropic glutamate receptor; GRIN: glutamate ionotropic receptor NMDA type; LRRTM: leucine-rich repeat transmembrane neuronal protein.

Receivers of postsynaptic glutamate signaling were inferred by expression of both ionotropic (*GRIA*, *GRIK*, *GRN*) and metabotropic (*GRM*) glutamate receptors (Fig 6c)^46^. As expected, neurons were the most prominent recipients of glutamate signaling during adulthood. Inferred neurexin-cerebellin to *GluD1* communications, which have been implicated in activity-dependent strengthening of glutamatergic synapses, were most prominent during infancy, consistent with the establishment of synapses and myelin during this period. By adulthood, inferred communication shifts to more mature synaptic interactions including both ionotropic (*GRIA*, *GRIK*, *GRN*) and metabotropic (*GRM*) glutamate receptors (Fig. 5c).

**Fig. 6.**
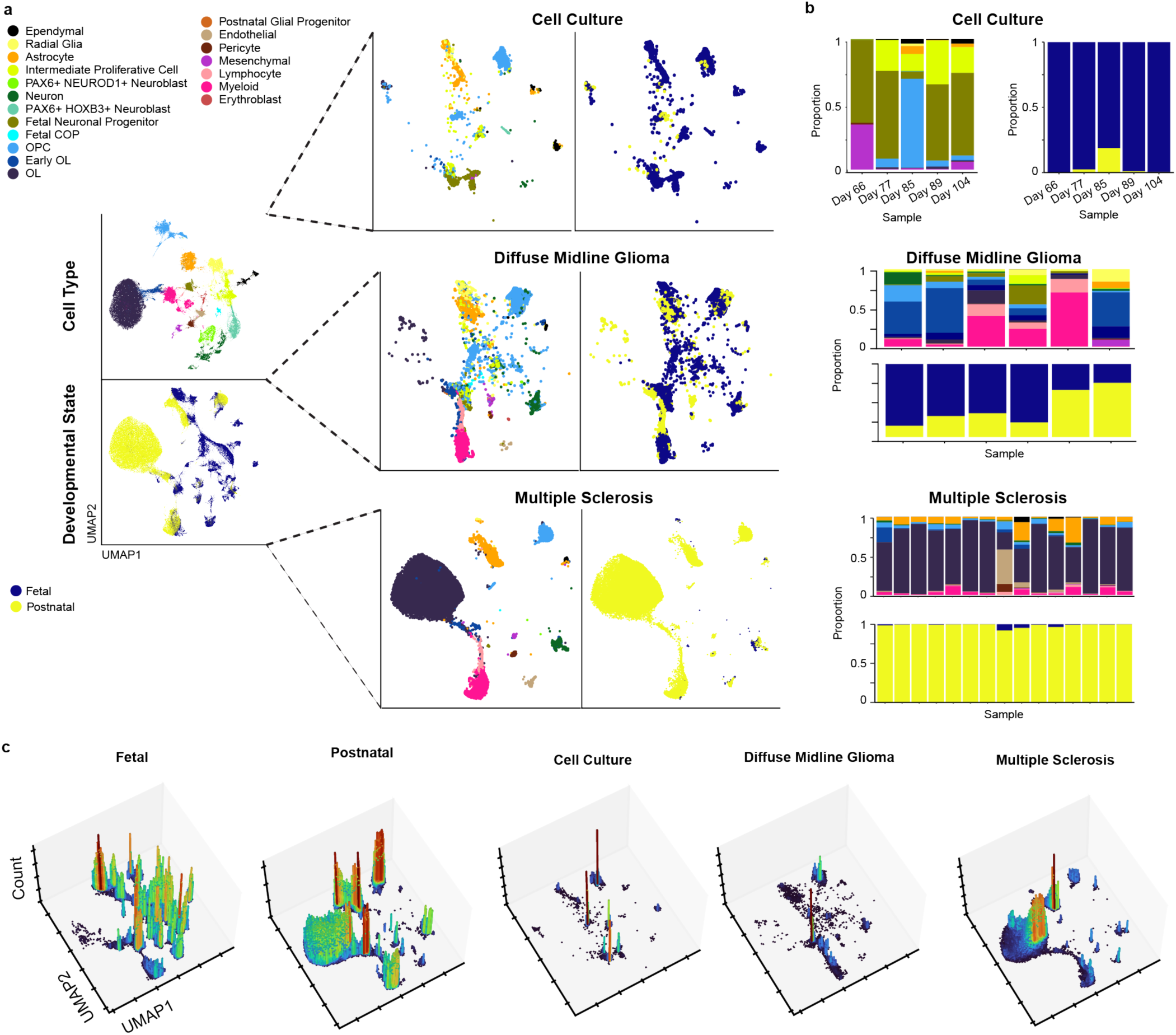
PEDIA-BRAIN provides context for vitro differentiation and disease states. Anchor-based mapping of *in vitro* differentiated pluripotent cells, pediatric high-grade gliomas, and multiple sclerosis white matter provides insight into research tools and disease processes. **a,** Comparison of *in vitro* differentiated oligodendrocyte lineage cells after 66, 77, 85, 89, and 104 days in culture (Top), pontine diffuse midline gliomas (n=6, Middle), and postmortem white matter from multiple sclerosis lesions (n=15, Bottom). **b,** Bar plots showing cell type proportions for cultured oligodendrocyte lineage cells (Top), pontine diffuse midline glioma (Middle), and multiple sclerosis white matter (Bottom). **c,** 3D UMAP of fetal, postnatal, cultured OPCs, multiple sclerosis, and pediatric glioma showing count of cells assigned to each UMAP coordinate. X-axis: RNA-UMAP-1, Y-axis: RNA-UMAP-2, Z-axis: Count of cells.

A critical period was noted for reinforcement of glutamate signaling between neurons and oligodendrocyte lineage clusters during the period from infancy to adolescence, via expression of the *GluD1/2* receptors involved in the neurexin-cerebellin transsynaptic strengthening pathway. Expression of both post-synaptic glutamate receptor subunits and genes involved in long term potentiation were observed during this critical period and were greatly diminished in adult OPCs (Fig. 5d). To explore the trajectory of glutamatergic synapses in the ventral pons white matter, the axial sections of the ventral pons from C57BL6 mice were stained for PSD95, a marker of the postsynaptic density in glutamatergic synapses, and PDGFRA, a marker of OPCs. While diffuse PSD95+ puncta were identified around PDGFRA+ OPCs during P2 and P8, by P24, when myelination of the corticospinal tract is complete, PSD95+ puncta were restricted to the actively myelinating transverse pontocerebellar fibers.

### PEDIA-BRAIN provides context for *in vitro* differentiation and disease states

Production of glia from pluripotent stem cells requires prolonged culture to induce the “gliogenic switch” from neurogenesis, which occurs *in utero,* to gliogenesis which occurs throughout life^16, 47^. To explore whether existing protocols recapitulate direct fetal trajectories or postnatal pathways that generate oligodendrocytes from homeostatic postnatal OPCs, 8,955 single cells from a published dataset of *PDGFRA+* OPCs generated from human pluripotent stem cells (GSE146373) were compared to the encyclopedia. While 88.9% of cultured cells mapped to fetal populations, prolonged *in vitro* differentiation successfully produced cells with postnatal signatures (Fig. 6a-b, Top, 6c)^48^.

To compare pontine diffuse midline gliomas to the normal developing pons, 25,508 single cell transcriptomes from 6 H3K27M-altered pontine diffuse midline gliomas were compared to our encyclopedia. Diverse fetal progenitor states were identified corresponding to radial glia, neuroblasts, and oligodendrocyte progenitor cells (Fig. 6a-b, Middle, 6c). A lack of cells with the transcriptional signature of mature oligodendrocytes was a unifying feature (55,497:85,416 OL:total in control postnatal vs 79:25,508 in DMGs, χ^2^ =32,852, p<0.00001). The ratio of early to mature oligodendrocytes was significantly higher in DMGs relative to other postnatal tissues, consistent with a differentiation block (7.6:1 early:mature ratio in DMG versus 1:17.2 in control postnatal, χ^2^ =7687.3, p<0.00001). While the ratio of radial glia to astrocytes was also skewed in DMGs compared to normal postnatal pons (1:1.21 RG:astrocyte ratio in DMG vs 1:71.6 in control postnatal, χ^2^ =3535.5, p<0.00001), the differentiation block appears to be uniquely severe for oligodendrocytes.

To explore the cellular diversity of multiple sclerosis, a disease that predominantly arises during adulthood, 52,856 single cells from the white matter of 15 human donors with multiple sclerosis (ages 35-68 years, GSE231858) were compared to the developmental encyclopedia of pons white matter (Figure 6a-b, Lower, 6c)^26^. Consistent with the age of donors, 95.5% (52,605/52,856) of MS white matter-derived cells mapped to postnatal cell types. The ratio of *RBFOX1*:*OPALIN* oligodendrocytes was skewed towards *OPALIN* in MS relative to control adult pons (22465:20036 in multiple sclerosis vs 10234:2509 in adult normal pons, χ^2^ =20,257, p<0.00001).

## Conclusions

We present a single nuclei-based encyclopedia of gene expression and chromatin accessibility in the human pons spanning the first four decades of life, as a resource for investigators studying disease processes in the pons. The full pons dataset and future datasets from ongoing projects targeting other brain regions can be accessed at https://pediabrain.nchgenomics.org. A searchable browser is provided to explore genes and chromatin peaks of interest with the capacity to output publication-quality figures. Links are provided to access data in raw and processed form along with code to map newly generated datasets to the encyclopedia.

The white matter of the ventral pons is an eloquent conduit for signals between brain regions and the body. Despite being a common site for pediatric gliomas and adult multiple sclerosis, our understanding of human pons white matter development is limited to morphometric and radiographic studies^4, 5^. The extent to which our increasingly sophisticated *in vitro* models recapitulate human disease lacks a reference for normal postnatal development. Similarly, our capacity to distinguish features of molecularly distinct disease states from normal developmental trajectories is hampered by the lack of a reference for normal postnatal development. PEDIA-BRAIN address this gap by creating a public reference for the study of developmentally timed brain disease. Although the work here focuses on the pons, our approach is applicable to other brain regions that exhibit developmental disease susceptibility.

At the completion of neurogenesis *in utero*, CNS progenitors undergo a neurogenic to gliogenic switch^16^. The low level of gliogenesis after infancy cannot account for the capacity of the brain to learn, however, postnatal brain development is likely reliant on epigenetically controlled changes in gene expression. As the narrow window of presentation for pontine H3K27M-altered DMGs points to a unique vulnerability of the pediatric pons to perturbations of chromatin regulation, we sought to generate a single nuclei atlas of gene expression and chromatin dynamics in the normal human pons from the first trimester to the completion of myelination in early adulthood.

Initial myelination of the pons occurs during infant and toddler years and is achieved by the generation of myelinating oligodendrocytes from OPCs^4^. While oligodendrocyte committed populations are identified in the fetal hindbrain, both the YAP1/SOX10+ committed oligodendrocyte and the fetal OPC population are enriched for regulatory factor X (RFX) transcription factor motifs while the homeostatic population of postnatal OPCs is enriched for TCF factors. Consistent with reports that oligodendrocytes may be generated directly from radial glia, unsupervised weighted nearest neighbor analysis indicates that both states converge on early oligodendrocytes^47, 49^.

Neuron-OPC synapses have been identified as a mechanism by which active neuronal circuits control OPC proliferation and direct their differentiation into myelinating oligodendrocytes^41^. Ablation of this “activity-dependent myelination” inhibits learning. Here we find that OPCs are highly enriched for synaptic processes relative to other oligodendrocyte lineage cells. OPC expression of organizers and receptors for glutamatergic synapses peaked during infancy and childhood and declined precipitously in adults. As neuronal firing similarly drives gliomas at neuron-glioma synapses, the striking correlation between the expression of synaptic plasticity genes in OPCs and the window of susceptibility to pontine DMGs represents a potential explanation for the predominance of pontine DMGs in children and warrants exploration in animal models^50^.

The classical function of oligodendrocytes in the CNS is facilitation of saltatory conduction; however, knockout studies have identified vital roles for oligodendrocytes in the provision of metabolic support for axons. Although several lines of evidence support both regional and grey-white specific functions, it remains unclear whether all oligodendrocytes perform these functions or whether these roles are fulfilled by subspecialized populations. Here we present a continuum of oligodendrocyte states defined by expression of the evolutionarily conserved mammalian myelin gene *OPALIN* or the RNA-binding protein *RBFOX1*, which is implicated in developmental delay and psychiatric disorders^44, 51^.

Our study is subject to sampling bias due to limited availability of human brainstem tissues. Additionally, the ventral pons is 50% white matter containing traversing axon bundles from distant sites with few local neuron cell bodies. Thus, the diversity of synaptic communications from the diverse neuronal populations that project to the ventral pons cannot be fully captured using assays that probe <10,000 cells and further studies using spatial transcriptomics are warranted to expand our understanding of synapse development in the pons white matter.

Our primary objective was to establish a benchmark of normal pons development and improve our understanding of developmental disease susceptibility. To illustrate the utility of the encyclopedia for evaluation of *in vitro* models and exploration of clinically relevant disease states, single cell transcriptomes from *in vitro* differentiated oligodendrocytes, pontine diffuse midline gliomas, and multiple sclerosis were compared to the encyclopedia. H3K27M-altered diffuse midline gliomas contained cells analogous to fetal progenitor states and early oligodendrocytes with a profound depletion of mature oligodendrocytes, consistent with a differentiation block. In contrast, samples with multiple sclerosis, a demyelinating disorder, exhibited a subtype-specific depletion of *RBFOX1+* oligodendrocytes enriched for metabolic processes and an increase in the fraction of cycling *OPALIN*+ oligodendrocytes that may represent ongoing remyelination.

Currently, validation of *in vitro* and *in vivo* disease models is based on conclusions drawn from cell types defined by a handful of canonical makers. Although bulk sequencing technologies have long provided insights into the gene expression and chromatin dynamics of cells that expand clonally in culture such as stem cells and tumors, single nuclei sequencing presents an opportunity to probe the diversity of differentiated cells that make up tissues in health and disease. Critically, the comparison of *in vitro* differentiated human oligodendrocyte progenitor cells to our encyclopedia of normal development permits an unbiased assessment of the degree to which *in vitro* models capture the complex interactions of the postnatal human brain. While bulk sequencing technologies are biased towards identification of changes driven by abundant cell types, profiling of single nuclei permits the identification of specific cell types that are depleted in disease states such our observation that DMGs lack mature oligodendrocytes and the *RBFOX1/OPALIN* ratio is altered in MS. Though this work focused on the pons, the methodology may be readily applied to the investigation of disease states affecting other brain regions or organ systems to provide further insights into human disease.

## Methods

### Animals and ethical statement

All procedures involving live animals were conducted in accordance with local regulations in an AAALAC-accredited vivarium. Procedures were approved by our Institutional Animal Care and Use Committee (IACUC: AR24-00104) and were designed to prioritize animal welfare by minimizing unnecessary pain and suffering.

### Human participants

#### Control Human Pons

Fresh frozen human pons tissue from donor autopsy specimens were obtained from the NIH NeuroBioBank (University of Maryland Brain and Tissue Bank) from the pool of specimens listed as “no clinical brain diagnosis” who were deceased from a “natural” cause of death. Donors aged <3 years of age were considered “infant and toddler” (n=4), >3-11 years of age were considered “child” (n=5), 13-14 years of age were considered adolescent (n=4), and >18 years of age were considered adult (n=4). Adult samples were selected from donors aged 31-33 as this age corresponds to the completion of myelination in the adult brain, overlaps with the age of onset for multiple sclerosis, and is prior to the onset of most non-immune neurodegenerative diseases. Socioeconomic status is not available from this de-identified biobank. Although an effort was made to obtain a balanced distribution of sexes, due to sample size and availability it was not feasible to sample a broad distribution of ethnicities.

#### DMG

Droplet based single cell RNA-sequencing using the 10X Genomics was performed on a previously unreported DMG sample obtained from stereotactic biopsy of a 6 year old male with a H3.3^K27M^-altered diffuse midline glioma at the time of progression.

| Sample ID | Age | Sex | Ethnicity | Condition | umbn* | GEO |
| --- | --- | --- | --- | --- | --- | --- |
| SC079_MW_1550_pons_s7 | 0 | Male | White | Normal | 1550 | GSE287571 |
| SC076_MW_771_pons_s7 | 2 | Male | Black or African-American | Normal | 771 | GSE287571 |
| SC075_MW_1275_pons_s7 | 2 | Female | Black or African-American | Normal | 1275 | GSE287571 |
| SC081_MW_1864_pons_sec7 | 2 | Female | White | Normal | 1864 | GSE287571 |
| SC089_MW_1499_pons_sec9 | 4 | Female | Asian | Normal | 1499 | GSE287571 |
| SC077_MW_4327_pons_S3 | 5 | Female | Black or African-American | Normal | 4327 | GSE287571 |
| SC088_MW_4332_pons_sec9 | 5 | Male | Black or African-American | Normal | 4332 | GSE287571 |
| SC082_MW_5925_pons_sec7 | 7 | Female | Black or African-American | Normal | 5925 | GSE287571 |
| SC084_MW_1860_pons_sec7 | 8 | Male | White | Normal | 1860 | GSE287571 |
| SC085_MW_1740_pons_sec7 | 13 | Female | Black or African-American | Normal | 1740 | GSE287571 |
| SC090_MW_4925_pons_sec7 | 13 | Male | Black or African-American | Normal | 4925 | GSE287571 |
| SC087_MW_4722_pons_sec7 | 14 | Male | White | Normal | 4722 | GSE287571 |
| SC080_MW_5309_pons_s7 | 14 | Female | White | Normal | 5309 | GSE287571 |
| SC128_MW_1538_pons_s5 | 32 | Male | Black or African-American | Normal | 1538 | GSE287571 |
| SC127_MW_1229_pons_s7 | 31 | Female | White | Normal | 1229 | GSE287571 |
| SC126_MW_1136_pons_s5 | 33 | Female | White | Normal | 1136 | GSE287571 |
| SC129_MW_4593_pons_s5 | 33 | Male | White | Normal | 4593 | GSE287571 |
| JL12_T | 6 | Male | White | Progressive H3K27M DMG |  | GSE287571 |

\*Refers to NIH NeuroBioBank (University of Maryland Brain and Tissue Bank) sample tracking number.

### Cell lines and reagents

No cell lines were used for this work.

Cell lines and reagents No cell lines were used for this work.
| Reagent | Vendor | Catalogue | RRID |
| --- | --- | --- | --- |
| Trizma Hydrochloride Solution, pH 7.4 | Sigma-Aldrich | Cat#T2194 |  |
| Sodium Chloride Solution, 5 M | Sigma-Aldrich | Cat#59222C |  |
| Magnesium Chloride Solution, 1M | Sigma-Aldrich | Cat#M1028 |  |
| IGEPAL CA-630 | Sigma-Aldrich | Cat#I8896 |  |
| Sigma Protector RNase inhibitor | Sigma-Aldrich | Cat#RNAINH-RO |  |
| DTT | Sigma-Aldrich | Cat#646563 |  |
| 7-AAD Ready Made Solution | Sigma-Aldrich | Cat#SML1633-1ML |  |
| 10% Tween 20 | Bio-Rad | Cat#1662404 |  |
| MACS BSA Stock Solution | Miltenyi-Biotec | Cat#130-091-376 |  |
| 1X Phosphate-Buffered Saline, pH 7.4 | Corning | Cat#21-040-CV |  |
| SPRIselect for Size Selection | Beckman-Coulter | Cat#B23319 |  |
| Buffer EB | Qiagen | Cat#19086 |  |
| Low TE Buffer (10 mM Tris-HCl pH 8.0, 0.1 mM EDTA) | Thermo Fisher Scientific | Cat#12090-015 |  |
| Glycerin (glycerol), 50% (v/v) Aqueous Solution | Ricca Chemical Company | Cat#3290-32 |  |
| Ethanol, Pure (200 Proof, anhydrous) | Millipore Sigma | Cat#E7023-500ML |  |
| Critical commercial assays |  |  |  |
| Chromium Next GEM Single Cell Multiome ATAC + Gene Expression Reagent Bundle, 16 rxns | 10X Genomics | Cat#PN-1000283 |  |
| Chromium Next GEM Chip J Single Cell Kit, 16 rxns | 10X Genomics | Cat#PN-1000230 |  |
| Single Index Kit N Set A, 96 rxns | 10X Genomics | Cat#PN-1000212 |  |
| Dual Index Kit TT Set A, 96 rxns | 10X Genomics | Cat#PN-1000215 |  |
| Nuclei Isolation from Complex Tissues for Single Cell Multiome ATAC + Gene Expression Sequencing | 10X Genomics | Cat#CG000375 Rev C |  |
| High Sensitivity D1000 ScreenTape/Reagents | Agilent | #067-5584/5067-5585 |  |
| High Sensitivity D5000 ScreenTape/Reagents | Agilent | #5067-5592/5067-5593 |  |
| Qubit dsDNA HS Assay Kit | Thermo Fisher Scientific | #Q32854 |  |

| **Equipment** |  |  |  |
| --- | --- | --- | --- |
| Bigfoot Spectral Cell Sorter | Thermo Fisher Scientific | #PL00304 |  |
| Chromium X | 10X Genomics | #PN-1000331 | RRID:SCR_024537 |
| BZ-X810 microscope | Keyence | #BZ-X810 | RRID:SCR_025160 |
| **Immunostaining Reagents** |  |  |  |
| OCT | Electron Microscopy Sciences | #62550-01 |  |
| Triton X | Sigma Aldrich | #X100-1L |  |
| Non-specific Donkey Serum | Sigma Aldrich | #9663-10mL |  |
| TrueBlack Lipofuscin Autofluorescence Quencher | Biotium | #23011 |  |
| DAPI ProLong Gold | Invitrogen | #P36931 |  |
| **Antibodies** |  |  |  |
| Rabbit anti-RBFOX1 (1:500) | Millipore Sigma | #HPA040809 | RRID:AB_10796228) |
| Guinea pig anti-Opalin (1:500) | Synaptic Systems | #457 005 | RRID:AB_2905562 |
| Guinea pig anti-SOX10 (1:1000) | Synaptic Systems | #347 404 | RRID:AB_3662608 |
| Rabbit anti-S100A1 (1:500) | ThermoFisher | #PA1-932 | RRID:AB_2183210 |
| Mouse anti-GFP (1:500) | Invitrogen | #MA5-15256 | RRID:AB_10979281 |
| Rabbit anti-YAP1 (1:200) | Proteintech | #13584-1-AP | RRID:AB_2218915 |
| Chicken anti-MBP (1:1000) | ThermoFisher | #PA-10008 | RRID:AB_1077024 |
| Goat anti-mousePDGFRA (1:500) | R&D Systems | #AF1062 | RRID:AB_2236897 |
| Mouse anti-PSD95 (1:500) | Synaptic Systems | #124 011 | RRID:AB_10804286 |
| **Mouse Strains** |  |  |  |
| C57BL/6J | The Jackson Laboratory | #000664 | RRID:IMSF_JAX:000664 |
| B6.129X1-Gt(ROSA)26Sortm1(EYFP)Cos/J (R26-loxI-STOP-loxP-EYFP) | The Jackson Laboratory | #006148 | RRID:IMSR_JAX:006148 |
| B6;CBA-Tg(Sox10-cre)1Wdr/J | The Jackson Laboratory | #025807 | RRID:IMSR_JAX:025807 |

#### Antibody Validation

Antibodies used for this study were validated by manufacturers for indicated applications. Additional validation included identification of expected morphology, cellular localization, co-staining with expected markers, and inclusion of positive/negative tissue controls. For detection of YFP in Sox10-Cre;R26-loxl-STOP-loxP-EYPF fate mapping studies, anti-GFP was used to detect YFP due to significant overlap in protein sequences of GFP and YFP and limited available anti-YFP host species. To confirm sensitivity of anti-GFP for YFP, Sox10-Cre dams were crossed with R26-loxl-STOP-loxP-EYPF males. E18.5 pups were harvested and genotyped to identify pups carrying the SOX10-Cre P1 chromosome. Utility of anti-GFP for detection of YFP was confirmed via detection of nuclear SOX10 and cytoplasmic GFP/YFP in ventral pons white matter and adjacent exiting cranial nerves in pups with SOX10-Cre P1 chromosome (n=2) and absence of signal in littermates without the SOX10-cre P1 artificial chromosome (n=1).

#### Histology

Human fresh frozen tissues were embedded in OCT prior to sectioning. Postnatal mice were anesthetized and euthanized via transcardiac perfusion with 4% paraformaldehyde, post-fixed for 24 hours, sunk in 30% sucrose, and embedded in OCT for sectioning in an axial plane. For studies with E18.5 embryos, the dam was anesthetized then pups were harvested, sunk in sucrose, and embedded in OCT for sectioning in an axial plane. OCT-embedded tissues were sectioned at 20μm, equilibrated to room temperature, and rehydrated in 1x PBS for 5 minutes. Sections were placed in a boiling 10% citrate solution and cooled for 30 minutes before neutralizing with a 5-minute wash in 1x PBS. Human sections were permeabilized in 2% TritonX for 30 minutes at room temperature. Sections were blocked in blocking buffer (10% Non-Specific Donkey Serum, 0.1% TritonX) for 1 hour at room temperature. Primary antibodies were diluted in blocking buffer and conjugated at 4 C overnight for mouse tissues and 3 days for human tissues. Sections were washed in 1x PBS 3 times for 5-8 minutes. Secondary antibodies were diluted in blocking buffer and conjugated for 1 hour in the dark at room temperature. Sections were washed in 1x PBS 3 times for 5-8 minutes. Sections were quenched in 30x TrueBlack Lipofuscin Autofluorescence Quencher (Biotium Cat No. 23011) diluted to 1x in 70% ethanol for 30 seconds – 2 minutes. Excess TrueBlack was dabbed off, and slides were rinsed 3 times in 1x PBS for 2 minutes. Glass coverslips were mounted using DAPI ProLong Gold (Invitrogen Cat No. P36931) and cured overnight in a dark, well-ventilated space.

#### Fate Mapping of Sox10+ cells in Murine Embryonic Pons

SOX10-Cre dams (Jax #025807) carry a P1 artificial chromosome with the Sox10 promoter driving Cre and were bred to ROSA26-LSL-EYFP (Jax #006148). Expression of the transcription factor SOX10 in myelinating cells of the central and peripheral nervous system in carriers leads to expression of YFP in all subsequent progeny. After confirmation of genotype in pups, studies were performed comparing SOX10-Cre;ROSA26-LSL-YFP pups versus littermates without SOX10-Cre. Due to the age of pups, lack of secondary sex characteristics prevented determination of biologic sex.

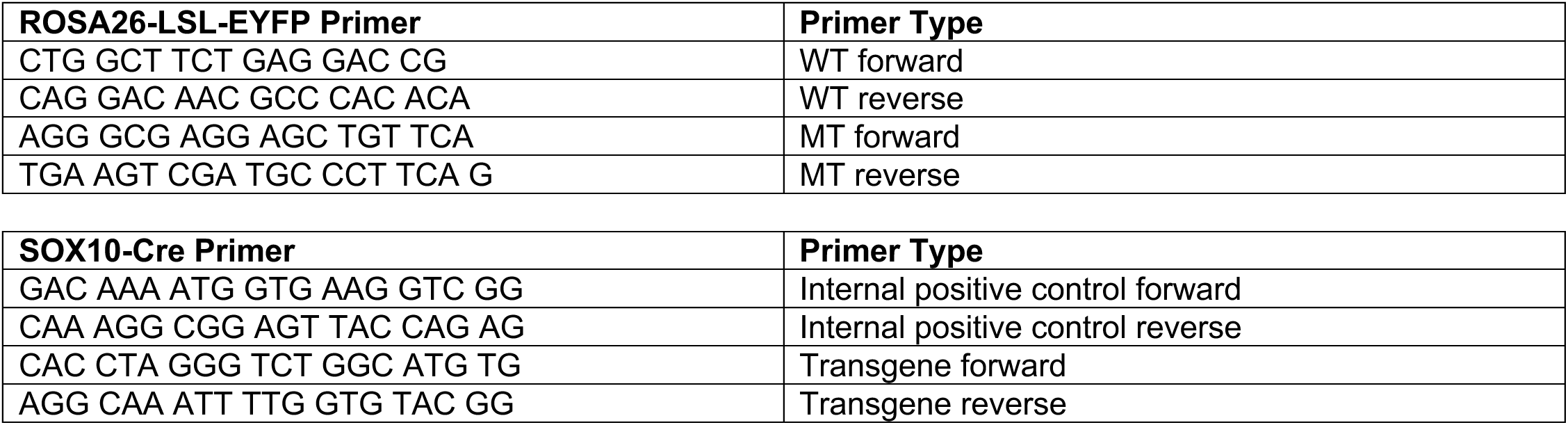

### Sample preprocessing and analysis

#### Nuclei isolation

Approximately 100 mg of frozen pons tissue was dissociated using the 10X Genomics protocol for Nuclei Isolation from Complex Tissues for Single Cell Multiome ATAC + Gene Expression Sequencing (CG000375 Rev C). The NP40 tissue lysis buffer consisted of the recommended alternative IGEPAL CA-630 at 0.1% instead of Nonidet P40 Substitute, and digitonin was not added to any of the buffers. A minimum of 500,000 7AAD+ nuclei were sorted using the ThermoFisher Scientific Bigfoot Spectral Cell Sorter. Nuclei permeabilization was then performed according to the manufacturer’s instructions.

#### Single nuclei RNA- and ATAC-seq library preparation and sequencing

Libraries were prepared from the permeabilized nuclei using the 10X Genomics Chromium Next GEM Single Cell Multiome ATAC + Gene Expression User Guide (CG000338 Rev F) according to the manufacturer’s instructions. Libraries were sequenced on an Illumina® NovaSeq6000 using two separate S2 Flow Cells v1.5 with a 100-cyle kit (≤2×50bp). Sequencing was performed using 50,000 paired-end reads per cell for each library. The read recipe for the dual-indexed Single Nuclei Gene Expression libraries was 50bp for Read 1, 8bp for the i7 Index Read, 24bp for the i5 Index Read, and 49bp for Read 2. For the dual-indexed Single Cell ATAC libraries, the read recipe was 28bp for Read 1, 10bp for the i7 Index Read, 10bp for the i5 Index Read, and 90bp for Read 2.

#### Fetal, pediatric, and adult single nucleus sequencing data preprocessing and integration

Raw base call (BCL) files were demultiplexed into FASTQ files, using 10x Genomics Cell Ranger ARC v2.0.0, and were subsequently aligned to a custom human hg38 reference and quantified using the Cell Ranger ARC “count” function^17^. Raw count matrices of 5 fetal samples were downloaded from the CATlas web browser (http://catlas.org/humanbraindev)^16^.

Count matrices from the 17 pediatric and adult samples and 5 fetal samples were loaded into R (v4.3.3) and merged using the Seurat (v5.1.0) and Signac (v1.14.0) packages^52^ ^53^. Quality control metrics were calculated according to standard Seurat/Signac workflows for gene expression and chromatin accessibility data. Doublets were identified by loading each sample count matrices into Python (v3.11.4) and calculating doublet score using the Scrublet package (v0.2.4)^54^.

Quality control was performed as follows to exclude low quality nuclei or duplicates prior to downstream analysis. Parameters were selected based on established practices and empirically based on the distribution of metrics to eliminate outliers. Nuclei with greater than 20% mitochondrial RNA were excluded. Nuclei with >30,000 RNA-seq molecules, >5,000 RNA-seq features, >80,000 ATAC molecules, >30,000 ATAC features, or a doublet score >0.25 were excluded from the final dataset as likely duplicates. After review of the distribution of quality-control parameters in our dataset, nuclei with poor quality ATAC-seq reads were excluded according to the following criteria: TSS enrichment score <1.5 reflecting decreased chromatin accessibility around transcription start sites relative to background, nucleosome signal >2.5 reflecting possible enrichment for nucleosome-bound fragments relative to mononucleosome, or blacklist ratio >0.05 reflecting poor quality sequencing. Pre- and post-QC density plots and number/fraction of cells filtered may be seen in Extended Data Fig. 1a-b. Many cells met more than one criterion for exclusion.

After QC, RNA-seq gene expression counts were merged, normalized, and scaled using SCTransform for identification of the top 50 principal components. Integration of the SCTransform counts was performed using the harmony package (v1.2.3) prior to principal component analysis and dimensionality reduction using uniform manifold approximately and projection (UMAP)^55, 56^.

ATAC-seq counts were analyzed in parallel. Normalization, variable feature identification and dimensionality reduction of ATAC counts were performed in Signac using Term Frequency-Inverse Document Frequency (TF-IDF) which prioritizes rare peaks as described by Stuart and Butler^28^. Merged chromatin accessibility data were integrated in harmony based on latent semantic indexing (LSI) coordinates followed by dimensionality reduction using UMAP.

Following separate dimensionality reduction using RNA and ATAC profiles, weighted nearest neighbor (WNN) integration was performed to achieve a joint low-dimensional embedding weighted by both RNA expression and chromatin accessibility.

#### Clustering and annotation of cell types

Clustering was performed in Seurat on harmonized gene expression counts with the Louvain algorithm from a resolution of 0.2 to 1.2 and visualized using Clustree to identify a resolution at which the clusters stabilized^57^. Cell types were annotated based on transcript expression of canonical marker genes in WNN clusters.

#### Differential gene expression (DEG) and Differentially Accessibly Peaks (DAP)

To reduce the impact of inter-individual variability on differential marker testing, raw RNA or ATAC counts for nuclei in each cluster were pseudobulked by sample using AggregateExpression() in Seurat. For each cell type pseudobulked values from each individual were used as inputs to DESeq2 for down differential gene expression testing between cell types, subtypes, and between age groups for astrocytes and OL lineages. For comparisons between cell types, transcripts with abs(log2(fold change))>1.5 corresponding to a 2.83-fold change and BH-corrected p-value<0.05 were considered significant. For comparisons between age groups for a given cell type, cell types with abs(log2(fold change))>1 corresponding to a 2-fold change and BH-corrected p-value<0.05 were considered significant. To identify differentially accessible peaks for each cluster, peak “counts” corresponding to the raw count of Tn5 insertion sites were pseudobulked by sample and DESeq2 was used to identify differentially expressed peaks. For comparisons between cell types, transcripts with abs(log2(fold change))>1.5 corresponding to a 2.83-fold change and BH-corrected p-value<0.05 were considered significant. For comparisons between age groups for a given cell type, peaks with abs(log2(fold change))>1 corresponding to a 2-fold change and BH-corrected p-value<0.05 were considered significant.

#### Motif enrichment, chromVar activity scoring, and transcription factor (TF) footprinting

Expressed transcription factors (TFs) for each cell type were defined as TFs that were expressed in at least 10% of nuclei in that cluster. Position weighted matrices for expressed motifs were obtained from the JASAPR 2024 database. Motif information was added to the Seurat object using AddMotifs followed by calculation of chromVar TF activity scores for each cell. chromVar scores are normalized across all cells. Overrepresentation analysis for expressed motifs was using FindMarkers in Signac. Regulatory TFs were identified as TFs with expression in >10% of cells in a cluster, log2(fold change)>1.5, and mean chromVar activity score >0.1. Footprinting of TFs was performed using Footprint() and PlotFootprint() in Signac.

#### Gene set enrichment analysis (GSEA)

GSEA of GO terms was performed with clusterProfiler (https://github.com/YuLab-SMU/clusterProfiler) with default parameters (adjusted maxGSSize = 1,000), and results were limited to enrichment in GO biological processes^58^. An FDR-adjusted *P* < 0.05 was considered significant.

#### Pseudotime analysis

A pseudotime trajectories of oligodendrocytes was built from RNA counts using the R package Monocle 3 (https://cole-trapnell-lab.github.io/monocle3/). PCA and UMAP were extracted from the Seurat object and Monocle3 “learn_graph” function was run^59^. The early OL cluster was set as the root cell. The “plot_cell_trajectory” function in Monocle3 was used to visualize the pseudotime trajectory across development.

#### Cell-cell communication network analysis

Inference of cell-cell communication probability for postnatal development was performed using the R package CellChat (v2) following the standard workflow as described in Jin, et al^60^. Cell types with >1000 cells were included for downstream analysis. Net communication probabilities were calculated as the sum of communication probabilities for each cell type and ranked in descending order to identify the top 5 pathways for each age group. To assess differences in cell communication pathways between age groups, an analysis of variance (ANOVA) test with Benjamini-Hochberg false discovery rate correction for multiple comparisons was applied. A corrected p-value of <0.05 was considered significant.

#### Preprocessing 10x Genomics based H3K27M-altered pontine gliomas and MS white matter

The following publicly available droplet-based single cell gene expression datasets were downloaded for mapping to the atlas: oligodendrocyte progenitor cells differentiated *in vitro* from human pluripotent stem lines (GSE146373), 15 multiple sclerosis samples (n=15, GSE231585), and pediatric diffuse midline glioma samples (n=5, GSE231860). An additional, previously unreported dataset was included from a post-treatment biopsy of a 6-year-old Caucasian male patient at our institution with an H3K27M-altered diffuse midline glioma was included (GSE287571) for a total of 6 pediatric diffuse midline gliomas. Raw FASTQs were processed as described previously then quality control filtered using the following criteria: number of RNA molecules less than 30,000, number of genes less than 6,000 (< 5,000 for the pediatric glioma samples), doublet score less than 0.25, and less than 20% mitochondrial RNA percentage.

#### Drop-seq based single-cell data preprocessing

FASTQ files were obtained from a publicly available single cell gene expression dataset generated by Drop-seq-based sequencing of human pluripotent stem cell-derived oligodendrocyte progenitor cells after 77, 89, and 104 days in culture (GSE146373). Quantified digital expression matrices (DEMs) were loaded into Seurat then analyzed as above from 10x-based single cell datasets. Gene expression data were quality control filtered using the following criteria: number of RNA molecules less than 30,000, number of genes less than 5,000, doublet score less than 0.25, and less than 20% mitochondrial RNA percentage.

#### Mapping of cultured oligodendrocyte lineage cells, multiple sclerosis, and pediatric glioma datasets to the atlas

Single cell transcriptomes from *in vitro* differentiated OPCs, pediatric high-grade gliomas, and multiple sclerosis were then mapped to the first 50 principal components of the reference pons atlas using the FindTransferAnchors function in Seurat. The resulting anchors were then used to map transcriptomes from the query datasets (*in vitro* differentiated OPCs, multiple sclerosis, pediatric high-grade gliomas) to the reference data set for assignment of cell types and developmental states using the MapQuery function in Seurat.

### Reporting summary

Further information on research design is available in the Nature Portfolio Reporting Summary linked to this article.

## Data availability

Data from these studies are available at pediabrain.nchgenomics.org. Metadata and RDS files of raw counts for each sample or group may be downloaded from this site along with processed data from the full encyclopedia.

Single-nuclei RNA-seq and ATAC-seq data have been deposited in the Gene Expression Omnibus and are publicly available as of the date of publication (GSE287571).

This paper analyzes existing, publicly available data from GSE146373, GSE287571, GSE231860, and GSE231585. Accession numbers for these datasets are listed in the key resources table. Raw data will be deposited by the time of publication at the National Institutes of Mental Health Neuroscience Data Archive (NDA) under collection C6003 PEDIA-BRAIN(Pediatric Encyclopedia of Development from Infancy to Adulthood - Brain Regions and Associated Integrative Neuroscience) with unrestricted access to the public.

## Code availability

All original code has been deposited at in GitHub and is publicly available as of the date of publication. All code necessary to reproduce our findings and examples for anchor-based mapping of outside datasets to the resource are available at github.com/Wedemeyer-Lab/PEDIA-BRAIN-Pons.

Any additional information required to reanalyze the data reported in this paper is available from the lead contact upon request.

## Acknowledgements

We are deeply grateful to James Fitch, Grant Lammi, and Adam Herman for management of the high performance computing environment.

Postnatal human pons samples were obtained from the NIH NeuroBioBank.

All sequencing was performed in the Nationwide Children’s Hospital Institute for Genomic Medicine’s Genomic Services Laboratory.

Funding was provided by a generous gift from Community Choice Financial, Inc. to the laboratory of Elaine Mardis, PhD.

## Author contributions

Conceptualization, M.A.W., K.E.M., and E.M.;

Methodology, M.A.W., E.M, K.E.M.;

Software Programming: T.D., G.S., A.V., K.L.S., A.H.W., B.K., K.Q, M.A.W;

Validation, M.A.W., T.D., A.R., K.K.;

Formal Analysis: G.S., T.D, A.V., M.A.W.;

Investigation: T.S., G.S., A.R., N.M., K.K.;

Resources: M.A.W., K.E.M, J.R.L., C.A.H., and E.M.;

Data Curation: A.V., G.S., T.D., K.S.;

Writing: Original Draft Preparation: M.A.W.;

Writing, Review & Editing: M.A.W, K.E.M, A.M.I., B.S., C.A.H., and E.M.;

Supervision: M.A.W., C.A.H., K.E.M., E.M.;

Project Administration: M.A.W., K.E.M;

Funding Acquisition: J.R.L., and E.M.

## Inclusion and ethics statement

The scope of this work using deceased, de-identified biobank specimens was determined to be “not human subjects research” by the Nationwide Children’s Hospital Institutional Review Board (STUDY00003079). The use of human tumor samples resected during neurosurgical procedures was approved by the Nationwide Children’s Hospital IRB (STUDY00003522, STUDY00000567, IRB16-00777). All methods were approved by our institutional biosafety committee (IBS00000693, IBS00000729).

---

**Extended Data Fig. 1.**
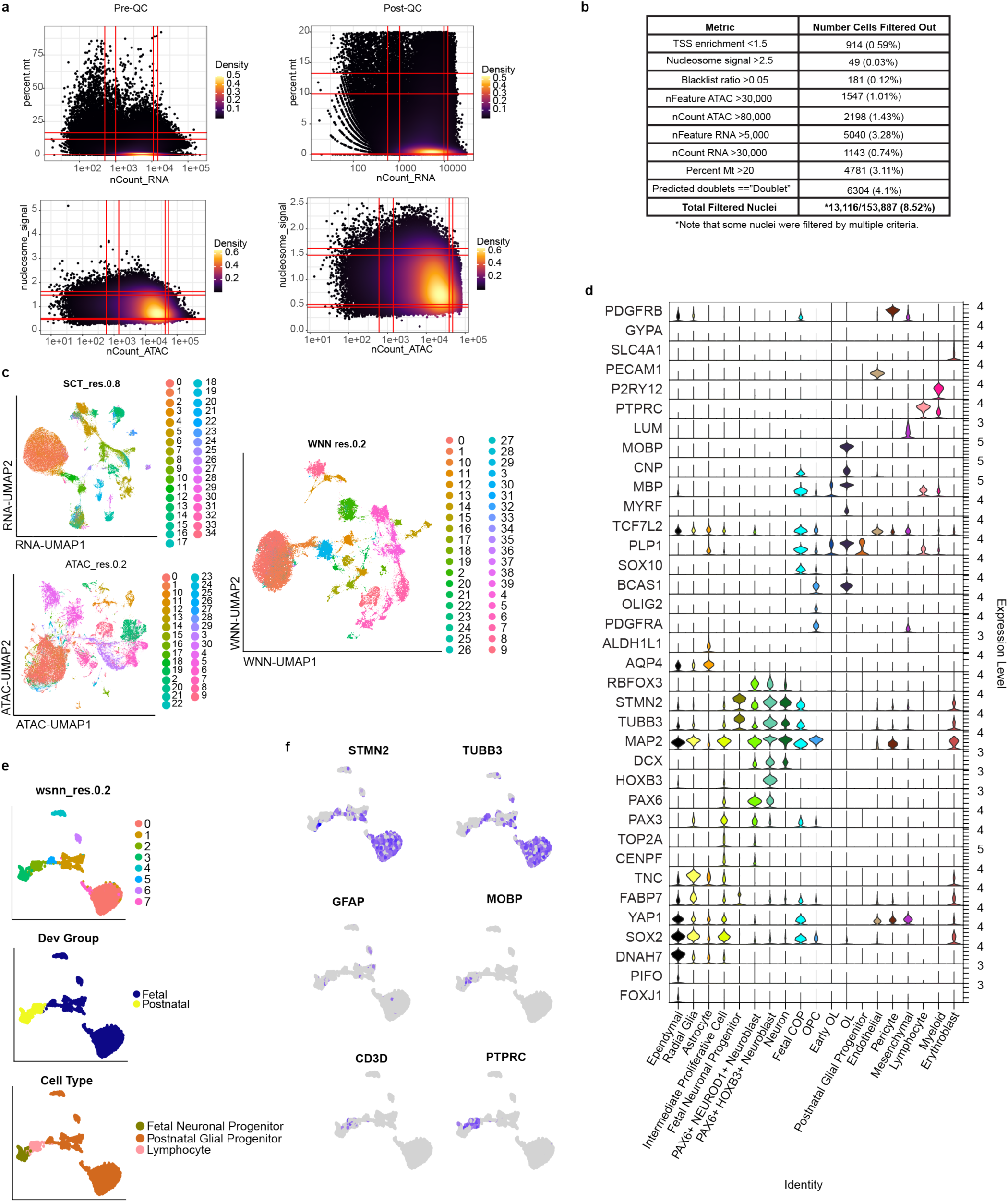
Combined single nuclei RNA+ATAC-sequencing identifies transcriptional signatures of cell types in the human pons from the first trimester to early adulthood. **a,** Density plot showing distribution of percent mitochondrial DNA, count of RNA, nucleosome signal, and number of ATAC counts before (left) and after (right) quality control to remove outliers. **b,** Table showing number of cells screened out by each parameter, with many meeting multiple criteria for removal. **c,** UMAP of encyclopedia clustered by RNA (res.0.8) showing 35 clusters, ATAC (res.0.2) showing 31 clusters, and WNN (res.0.2) showing 40 consensus clusters. **d,** Violin plot of canonical markers used for cell type assignment. **e,** Subclustering was performed on cluster 11 which was driven by high expression of lymphocyte genes (*PTPRC*, *CD3D*) in subcluster 2. Subclustered WNN UMAP grouped by WNN cluster, fetal vs postnatal, and assigned subtype. **f,** Feature plots of subclusters from cluster 11 showing a single cluster of lymphocytes (CD3D, PTPRC) while remaining clusters expressed genes broadly expressed in neuronal progenitor (*STMN2*, *TUBB3*) or glial (*GFAP*, *MOBP*) progenitor cell types.

**Extended Data Fig. 2.**
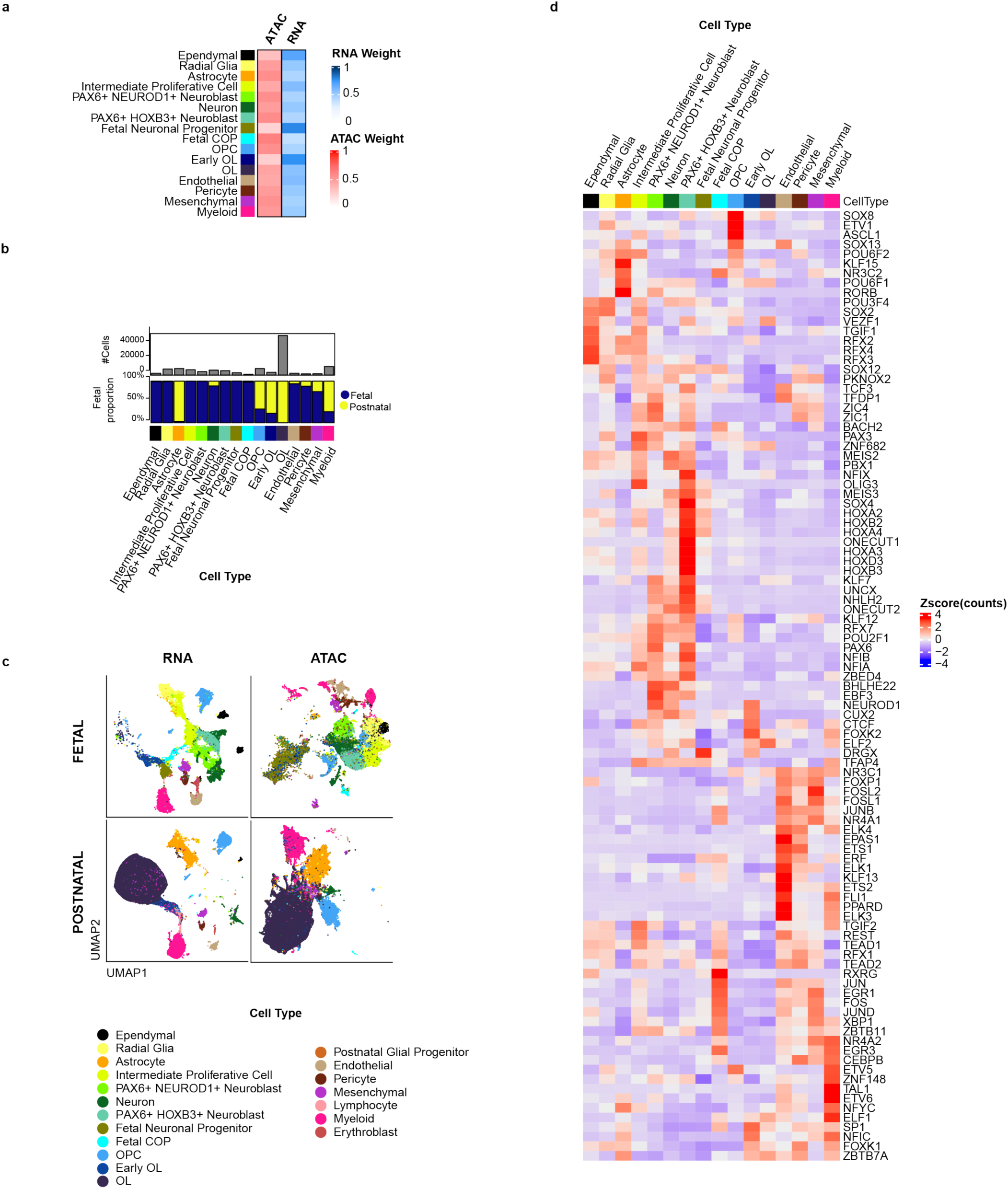
Integrated analysis of gene expression and chromatin accessibility reveals the cellular composition and transcription factor regulators of the developing pons. **a,** Heatmap showing contribution of RNA versus chromatin accessibility signatures to the weighted nearest neighbor signature for major cell types in the pons. **b,** Summary of cell counts and tissue of origin for clusters with >1000 cells. Note that several progenitor states are almost exclusively identified in fetal tissues. **c,** RNA (left) and ATAC (right) UMAPs for cells derived from fetal (top) versus postnatal tissues (right). **d,** Heatmap showing normalized RNA counts for the top 10 transcription factors whose motifs are significantly enriched in clusters with >1000 cells. Motif enrichment was performed on the top 5000 differentially accessible peaks in each cluster using motifs for TFs with expression in >10% of nuclei in at least one cluster. Motifs with abs(log2(fold change))>1.5 relative to background and BH-corrected p-value<0.05 were considered enriched.

**Extended Data Fig. 3.**
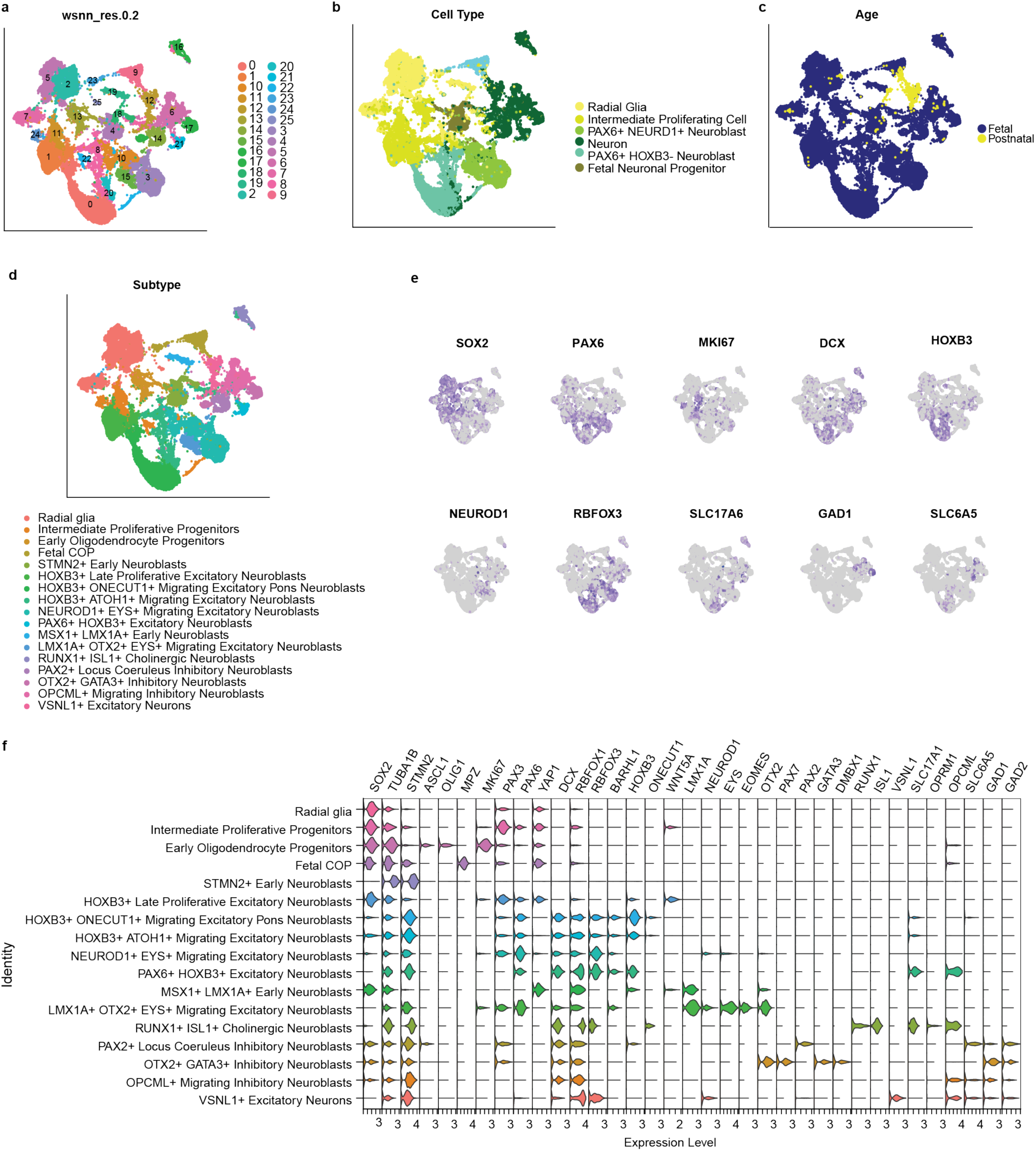
Developmental trajectory of radial glia and astrocytes. **a-c,** WNN UMAP of subclustered (res 0.2) radial glia, neuronal progenitor, and neuron clusters identified 26 sub-clusters. Cells are grouped by subcluster (**a**), cell type (**b**), and age group (**c**). **d,** UMAP of assigned subtype. **e,** Feature plots showing distribution of radial glia and intermediate progenitors (*SOX2*, *PAX6*, *MKI67*), migrating neuroblasts (*DCX*), hindbrain identify (*HOXB3*), maturing neurons (*NEUROD1*, *RBFOX3*), excitatory neurons (*SCL17A6/VGLUT2*), inhibitory neurons (*GAD1*, *GAD2*), and glycinergic neurons (*SLC6A6*, glycine transporter 2/norepinephrine receptor 1). **f**, Violin plot of features used to assign subtypes.

**Extended Data Fig. 4.**
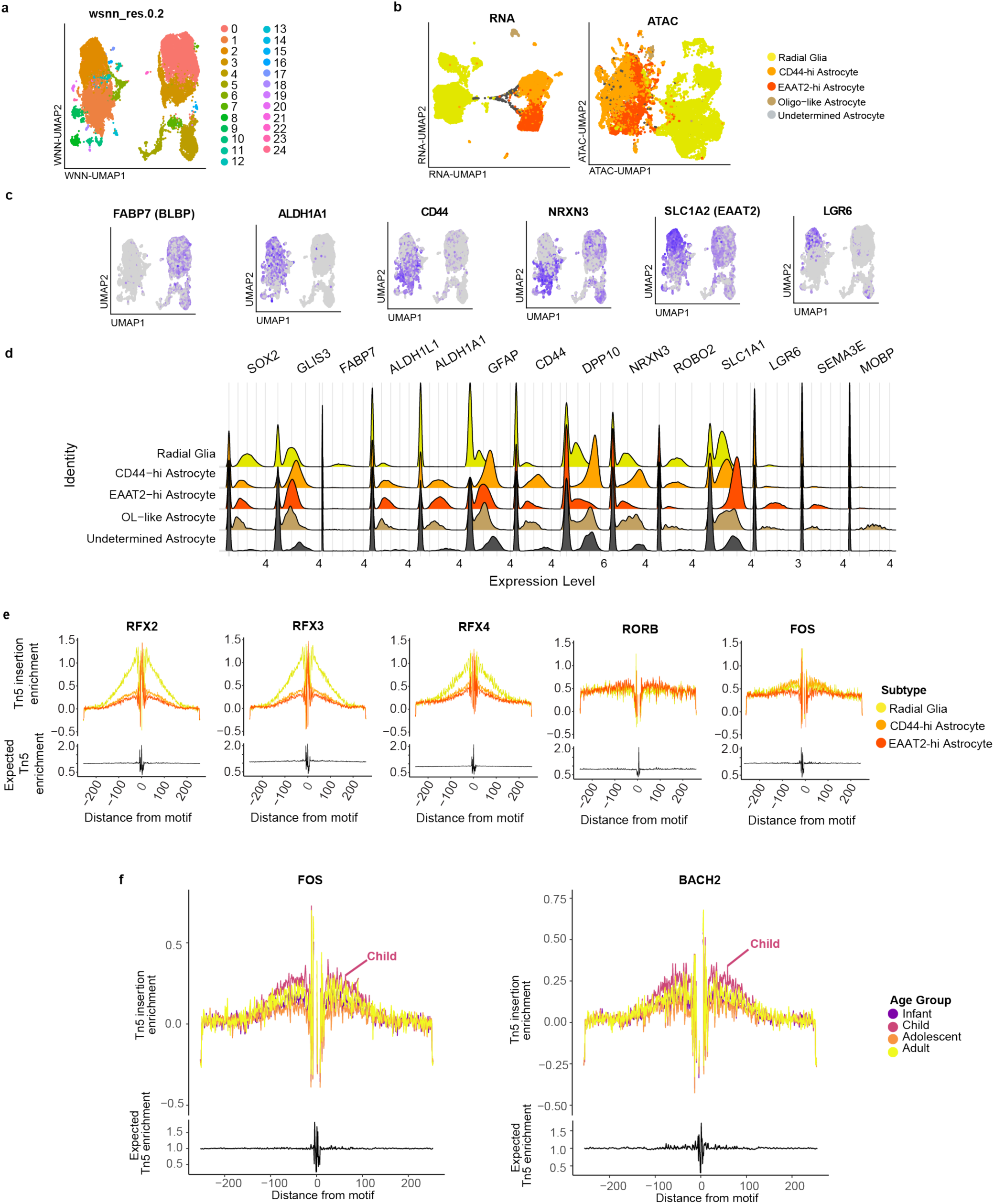
Gene expression and chromatin accessibility in radial glia and astrocytes. **a,** WNN UMAP of radial glia and astrocytes identified 25 sub-clusters. **b,** UMAP of radial glia and astrocytes subclustered by RNA (Left) and ATAC (Right) and grouped by assigned subtype. **c,** Feature plots of canonical markers used to classify subclusters shows increased expression of the stem cell markers *FABP7* in radial glia and *ALDH1A1* in postnatal astrocytes. Two major subtypes of astrocytes are distinguished by expression of *CD44/NRXN3* (*CD44-hi*) versus *SLC1A1(EAAT2)/(LGR6* (*EAAT2-hi*). **d,** Ridge plot of top genes that distinguish subtypes of radial glia and astrocytes. **e,** Footprint plots of transcription factors with significantly different motif enrichment radial glia versus astrocyte subtypes. A dip at the motif binding site (0) relative to bias-corrected Tn5 insertion probability is suggestive of protection from Tn5 insertion due to transcription factor occupancy. An increase in Tn5 signal from baseline in regions flanking a motif denotes increased insertion frequency and is characteristic of active regulatory regions with increased chromatin accessibility. **f,** Footprint plots for *FOS* and *BACH2* across postnatal development of *CD44-hi* astrocytes showing increased chromatin accessibility for both TFs during childhood.

**Extended Data Fig. 5.**
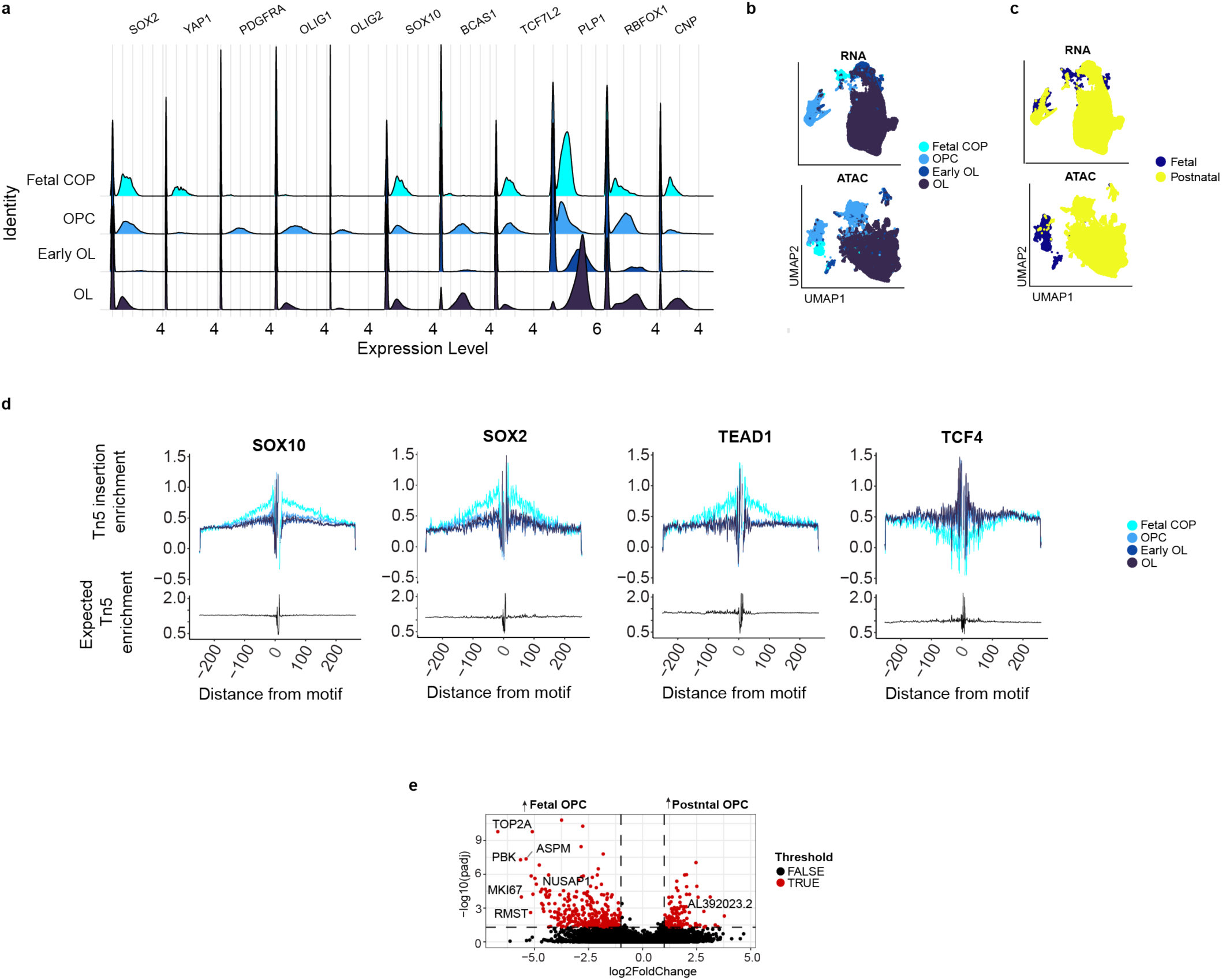
Gene expression and chromatin accessibility in oligodendrocyte lineage cells. **a,** Ridge plot of canonical markers used to classify oligodendrocyte lineage subclusters. Fetal COPs express radial glia markers (*SOX2*, *YAP1*) while OPCs express canonical oligodendrocyte lineage transcription factors *OLIG1/2*. **b-c,** RNA (top) and ATAC (bottom) UMAP of oligodendrocyte progenitor/precursor lineage subclusters grouped by subcluster (**b**) and tissue of origin (**c**). **d,** Footprint plots of transcription factors with significantly enriched motifs in each subtype relative to other oligodendrocyte lineage cell types. A dip at the motif binding site (0) relative to bias-corrected Tn5 insertion probability is suggestive of protection from Tn5 insertion due to transcription factor occupancy. An increase in Tn5 insertion frequency from bias-corrected baseline denotes increased insertion frequency in flanking regions and is characteristic of increased chromatin accessibility at active regulatory regions. **e,** Volcano plot of transcript expression in fetal vs. postnatal OPC. Red dots denote genes with BH-corrected p-value<0.05 and abs(log_2_(fold change)) >1.5.

**Extended Data Fig. 6.**
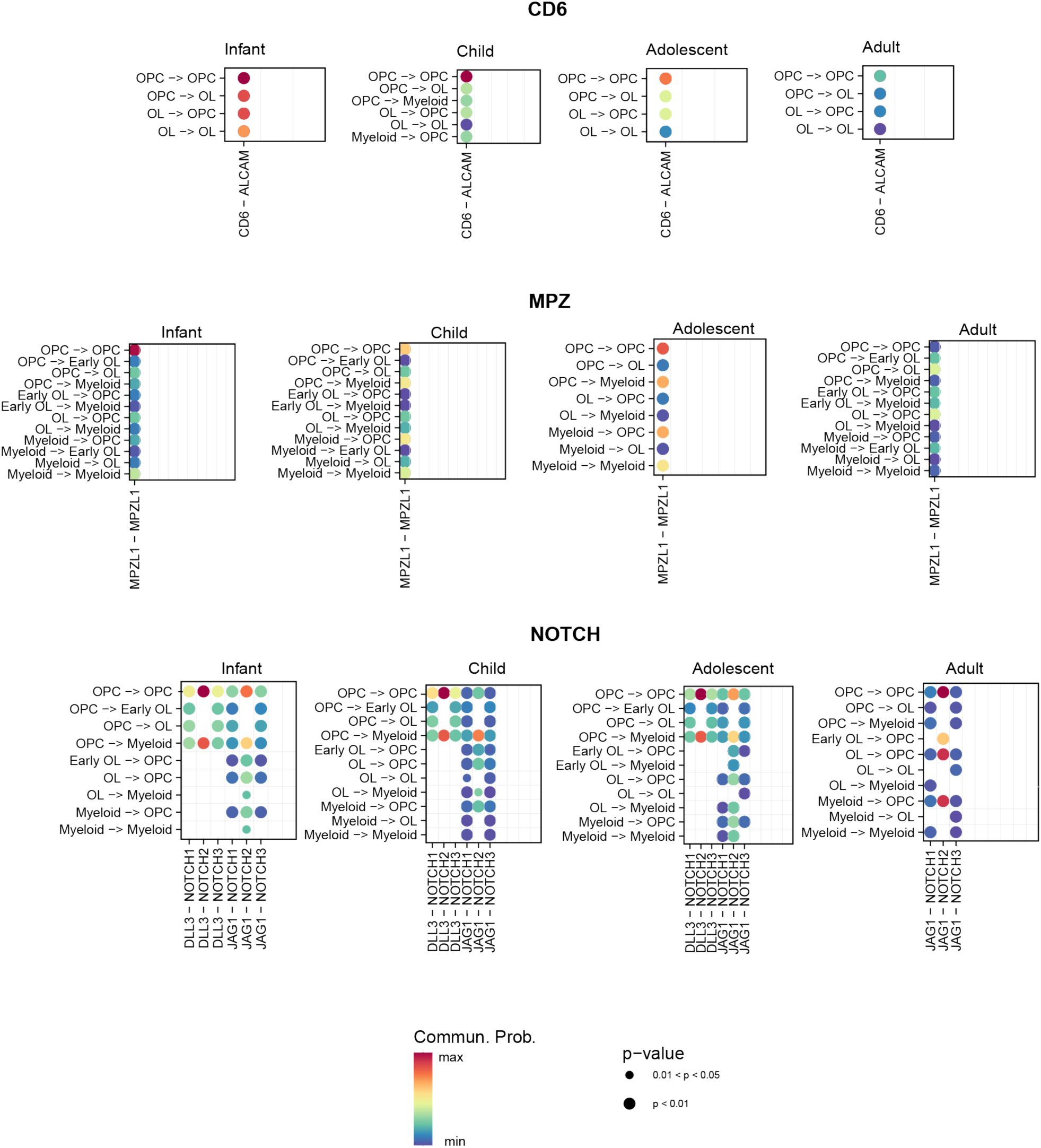
CD6, NOTCH, and MPZ pathways exhibit developmental shifts in receptor ligand signaling probability. Dot plot of communication probabilities for CD6 (Top), MPZ (Middle), and NOTCH (Bottom) signaling between postnatal cell types with >1000 cells. Dot colors represent inferred receptor-ligand communication probability while dot size represents the BH-corrected p-value for the inferred communication.

## Supplementary Tables

Supplementary Table 1: Differentially expressed genes for each cell type

Supplementary Table 2: Differentially accessible peaks for each cell type

Supplementary Table 3: Differentially enriched motifs for each cell type

Supplementary Table 4: Enriched biological processes in cell types and subtypes

Supplementary Table 5: Differentially expressed genes, accessible peaks, and enriched motifs in astrocyte and oligodendrocyte subtypes

Supplementary Table 6: Differentially expressed genes, accessible peaks and enriched motifs in postnatal astrocyte and oligodendrocyte subtypes across age groups

Supplementary Table 7: Cell-cell communication pathway probabilities

